# Background selection in recombining genomes and its consequences for the maintenance of variation in complex traits

**DOI:** 10.1101/2025.05.27.656342

**Authors:** Xinyi Li, Jeremy J Berg

## Abstract

Background selection (BGS)—the reduction of linked neutral diversity via the purging of deleterious mutations—is a pervasive force in genomic evolution. However, its impact on complex phenotypes remains poorly understood because classical theory treats fitness effects as fixed rather than emerging from a phenotype-to-fitness map. Here, we investigate the impact of BGS across three phenotypic selection frameworks: exponential directional, a liability threshold model, and stabilizing selection. First, we develop an effectively non-recombining block approximation for the site frequency spectrum (SFS) and show that this framework accurately describes the skew in the site frequency spectrum in the weak mutation regime typical of humans. Second, we show that phenotypic impacts of BGS depend on how selection is coupled across loci. In the liability threshold model, strong synergistic epistasis generates a global compensation mechanism—driven by tiny shifts in the mean phenotype— that propagates BGS effects to strongly selected variants otherwise immune to linked selection. This coupling reduces genetic variance across almost the entire effect-size distribution by an amount determined by a non-linear average of local effective population size reductions across the genome. Conversely, under stabilizing selection, BGS can counter-intuitively increase genetic variance. This occurs because BGS shifts strongly selected sites into the weakly selected underdominant regime where they persist at intermediate frequencies longer than in the equivalent directional selection model. Our results inform both longstanding evolutionary conversations regarding synergistic epistasis and efforts to model the impact of background selection on individual variants in recombining genomes.

**Significance:** Natural selection against deleterious mutations reshapes genetic variation through background selection, a process typically modeled as a local reduction in effective population size. We show that the consequences of this reduction for genetic architecture are mediated by the relationship between phenotype and fitness. Extending background selection theory to complex traits, we show that in models with strong synergistic epistasis, a global compensation mechanism propagates the effects of selection to strongly selected variants. Furthermore, we find that background selection can counter-intuitively increase genetic variation under stabilizing selection. Our work shows that the total impact of background selection on functional variation thus depends on the answers to longstanding questions about the ultimate sources of variance in fitness.

## Introduction

Natural selection acting on deleterious mutations reduces genetic variation at linked neutral sites, a process known as background selection (BGS; Charlesworth et al. 1993). In the decades since its initial description, BGS has become a standard component of population genetic null models. To a first approximation, BGS acts as a local reduction in the effective population size (*B* = *N*_*e*_*/N*), which reduces the expected number of segregating sites and increases the rate of genetic drift (Hudson and Kaplan 1995; Nordborg et al. 1996; McVicker et al. 2009). The magnitude of this reduction depends largely on the local density of functional sites relative to the recombination rate and varies substantially across species. For example, in humans, BGS effects are generally modest (*B* ≈ 0.6 in the most affected regions, with *B* ≈ 0.83 on average; Murphy et al. (2022) and Buffalo and Kern (2024)), whereas in the more compact *Drosophila* genome, the average reduction is much stronger (mean *B* ≈ 0.3; Comeron (2014) and Elyashiv et al. (2016)). The existence of broad-scale correlations between recombination rate and diversity across taxa suggests that these linked selection effects are a pervasive feature of genomic evolution across the tree of life (Corbett-Detig et al. 2015; Coop 2016; Buffalo 2021).

Despite this progress, our understanding of how BGS impacts the evolution of complex phenotypes remains limited. Empirical studies have suggested that BGS shapes the genetic architecture of traits. For example, Rockman et al. (2010) observed that in *C. elegans*, loci in gene-poor, high-recombination regions explain significantly more variance in gene expression than those in regions with higher gene density and lower recombination rate. Similarly, in humans complex traits, per-SNP heritability appears enriched in regions of strong background selection (Gazal et al. 2018; Pardiñas et al. 2018). However, theoretical work has largely been restricted to single-site models in which the fitness effects of individual loci are fixed (Charlesworth 1994; Stephan et al. 1999). However, assuming that the fitness effects of variants are largely mediated by their contributions to polygenic traits, their fitness effects are not fixed, but a emerge from the distribution of phenotypes in the population and the relationship between phenotype and fitness (Simons et al. 2018; Berg et al. 2025). Consequently, determining the full impact of background selection requires understanding how these local linkage effects interact with the global, polygenic constraints that ultimately generate fitness.

Even at the level of individual sites, the standard “reduced local *N*_*e*_” approximation provides an incomplete picture. While it accurately predicts the total heterozygosity and fixation rate, it fails to capture the distortion of the site frequency spectrum (SFS) relative to a simple population size reduction (Gordo et al. 2002). This distortion arises because BGS does not act instantaneously; alleles must persist long enough to “feel” the purging of linked deleterious backgrounds. This delay creates a skew in the SFS that can confound both demographic inference and inferences about the distribution of fitness effects (Johri et al. 2021). Prior work has provided an analytical description of this skew for non-recombining regions in the strong mutation regime (Cvijović et al. 2018) . Here, we develop an effectively non-recombining block approximation for the dynamics of BGS in recombining genomes, and use it describe the skew in the frequency spectrum for in the weak mutation limit, and show that is valid broadly in the parameter regime relevant to humans.

We then study the impact of BGS in three models of phenotypic mutation-selection-drift balance. We first contrast two models of directional selection: an exponential fitness model, where loci evolve independently, and a liability threshold model, where fitness effects are tightly coupled across loci. We show that in the threshold model, the system responds to BGS through a global compensation mechanism driven largely by the genomic regions experiencing the strongest background selection. Unlike simple local *N*_*e*_ rescaling—which impacts only weakly selected sites—this global mechanism propagates the effects of BGS to strongly selected variants that would otherwise be immune to linked selection. Finally, we consider a model of stabilizing selection, the ubiquity of which is motivated by both direct (Karn and Penrose 1951; Sanjak et al. 2018) and indirect (Smith 1983; Simons et al. 2022; Koch et al. 2024) lines of evidence. We find that due to the specific dynamics of underdominant selection maintained by stabilizing selection, BGS can, counter-intuitively, lead to an *increase* in the genetic variance of quantitative traits.

## Results

Our results are presented in three parts. First, we consider a model of BGS acting on a single focal site in a recombining background. We focus on modeling the skew in the frequency spectrum under moderate BGS strengths and resolving the distinct effects of BGS on alleles under weak versus strong direct selection. Second, we apply results from this single site analysis to models of polygenic directional selection. We contrast a ‘decoupled’ exponential fitness model, where BGS effects are entirely local, with a ‘coupled’ threshold model, where strong epistasis forces a global, compensatory response. Finally, we analyze a stabilizing selection model to demonstrate how a different form of epistasis leads to a distinct, local-only BGS dynamic, which can lead to increases in genetic variance under some conditions.

### Background selection in single site models

#### The frequency spectrum

##### Neutral alleles

We begin by considering a diploid population of size *N*, focusing on an individual site embedded in a recombining chromosome of length *M* . Adjacent sites recombine with rate *r*, and each produces deleterious mutations (with selection coefficient *s*_*b*_) at rate *ν*. We further assume selection against these mutations is strong relative to drift (i.e., 2*Ns*_*b*_ ≫ 1, ensuring they are purged and never fix), but weak relative to the chromosome’s total recombination rate (i.e., *s*_*b*_ ≪ *Mr*). For the moment, we suppose this site is neutral, but will relax this shortly.

Under these assumptions, the impact of BGS can be understood through an “effectively non-recombining linkage block” approximation (see Supplementary Text A; Neher et al. 2013; Good et al. 2014; Weissman and Hallatschek 2014). This model approximates the background as a single non-recombining block of characteristic length 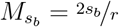, a scale set by the balance between the purging of deleterious mutations and the erosion of the haplotype by recombination. Because the expected frequency of a deleterious allele at any background site is *ν/s*_*b*_, this characteristic block carries an average of

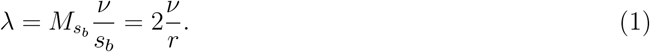

deleterious alleles (Eq. (A.4)).

While it is the burden of deleterious mutations within the characteristic length governs the classic BGS process, newly arising mutations also exert an influence in early generations. Immediately upon arising, the focal allele becomes associated with an average of *λ* additional deleterious mutations distributed over a physical distance much larger than the characteristic length-scale, an expectation that remains approximately constant for short times due to the balance between mutation and recombination. In Supplementary Text A, we develop a heuristic “shrinking block” approximation for the impact of this additional fitness drag, and describe how it differs from the analogous process of fitness decline in a non-recombining model.

We find that if mutation is weak relative to recombination (more precisely, if *λ* ≲ 1*/*2), then the characteristic haplotype block does not accumulate enough fitness drag to have an impact before selection can act against the existing background. Consequently, if a focal allele is eliminated by selection against a linked deleterious allele, it is almost certainly due to an allele that was already present when it arose, and so mutations outside the characteristic length-scale are irrelevant. Under these assumptions, the expected deleterious load on a haplotype block of length 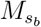 at frequency *q* is exponentially suppressed to 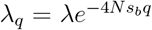 . To reach frequency *q*, a focal allele must therefore reside on a background that carries, on average, 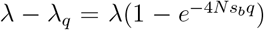 fewer deleterious mutations than a background chosen at random. This purging process reduces the number of alleles that persist to frequency *q* by a factor equal to the survival probability of the background:

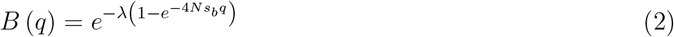

(see Eq. (A.26)), which converges to the standard “reduced effective population size” approximation, *B* = *e*^−*λ*^, for for 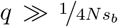 (Hudson and Kaplan 1995; Nordborg et al. 1996). Thus, in the weak mutation limit, the skew in the frequency spectrum mirrors the exponential suppression of frequencies among the deleterious alleles responsible for BGS (i.e. 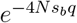). This approximation can be straight-forwardly extended to a distribution over *s*_*b*_ by averaging the exponential suppression term, giving the frequency-dependent reduction factor:

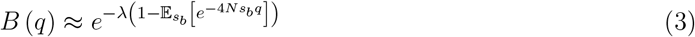

(see Eq. (A.27)). While the assumptions justifying these approximations are only formally satisfied in the limit as *λ* → 0, in practice we find that they provide a good fit for the *λ* ≲ 1*/*2 (see Figure S1 and Figure 1 below), which corresponds to *B* ≥ 0.6, encompassing roughly the full range of background selection effects observed in humans (Murphy et al. 2022; Buffalo and Kern 2024).

##### Directly selected alleles

Now, suppose the focal site itself produces alleles experiencing additive direct selection with coefficient *s*, and let 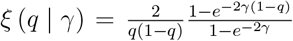 be the number of generations that such an allele spends at frequency *q* during its transit through the population, where *γ* = 2*Ns* is the scaled selection coefficient (Wright 1938; Bustamante et al. 2001).

When direct selection on the focal allele is weak relative to background selection (|*γ*| ≪ 2*Ns*_*b*_), it influences dynamics only at high frequencies 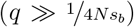, where the initial cull of alleles arising on deleterious backgrounds is complete (*B*(*q*) = *e*^−*λ*^). This convergence reflects a separation of timescales: because haplotypes carrying deleterious alleles are rapidly purged, any focal allele reaching high frequency must have arisen on a background that is free of deleterious mutations over the characteristic length-scale (Charlesworth 1994; Stephan et al. 1999). A new, weakly selected mutation is thus expected to spend

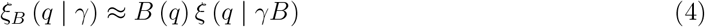

generations at frequency *q* (see Eq. (A.30)). Consequently, for deleterious alleles under direct selection weaker than the background intensity, BGS reduces sojourn times at low frequencies but increases them at high frequencies. Notably, this effect persists even for alleles that are strongly selected relative to drift (|*γ*| ≫ 1), provided they satisfy |*γ*| ≪ 2*Ns*_*b*_ (Figure 1B). For beneficial alleles, these effects compound, further suppressing the number of alleles that reach high frequencies (Figure S2).

Finally, when direct selection is strong (|*γ*| ≳ 2*Ns*_*b*_), direct and background selection act on similar timescales. The allele’s fate is determined by its own fitness effect before the background is purged, suggesting the alternative approximation

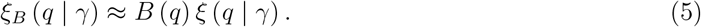

(Eq. (A.35)). In practice, however, these strongly selected alleles rarely reach the high frequencies where background selection takes effect (*q* ≫ 1*/*4*N s*_*b*_). Consequently, for the vast majority of variants in this regime, *B*(*q*) ≈ 1 and BGS is irrelevant. While we do observe some distortions in the frequency spectrum for *q* ≳ 1*/*4*N s*_*b*_ (Figure 1C), these rare variants contribute minimally to standing variation.

### Phenotypes under directional selection

#### Heterozygosity and fixation rates at individual sites

We now turn to models of mutation-selection-drift balance (MSDB) for polygenic phenotypes. We analyze these models using a mean-field approach, approximating the evolution of the phenotypic distribution by the average behavior of a single site. The key features of this behavior are captured by two quantities: an allele’s fixation probability and its expected lifetime contribution to heterozygosity.

As shown by Kimura (1969), these two quantities are intimately related, as the lifetime contribution to heterozygosity is:

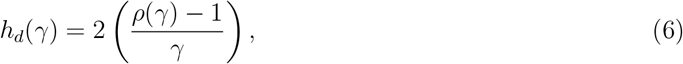

where 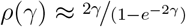 is the expected number of fixations per 2*N* new mutations, and the quantity *ρ*(*γ*) − 1 is the “selective fixation rate” (the number of fixations driven or prevented by selection). To model the impact of background selection, we employ the standard reduced effective size approximation (*B* = *e*^−*λ*^). While equation (4) breaks down for *B* ≲ 0.6, this approximation remains accurate for the heterozygosity provided that 2*NBs*_*b*_ ≫ 1—a condition that ensures local effective size reductions are not severe enough to push sites into the interference selection regime (Good et al. 2014). This means that our phenotypic results should apply in the presence of much greater effective size reductions than our simple approximation for the skew.

Applying this approximation, the reduction in the lifetime contribution to heterozygosity due to BGS is precisely equal to the effect on the selective fixation rate:

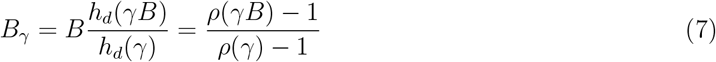

where the factor of *B* represents the probability of surviving the initial low-frequency cull, and *h*_*d*_(*γB*)*/h*_*d*_(*γ*) represents the effect on the lifetime contribution to heterozygosity conditional on surviving this cull (Figure 2A). Notably, this relationship holds even for strongly selected sites (*γ* ≪ −1) where the physical mechanism differs. In the regime where direct selection is strong relative to drift but weak relative to background selection (2*Ns*_*b*_ ≫ |*γ*| ≫ 1; Figure 1B), BGS significantly distorts the frequency spectrum: it reduces the number of alleles that survive the initial purge, but allows the survivors to drift to higher frequencies due to weakened effective selection. These opposing effects cancel out, leaving the total heterozygosity unchanged (*B*_*γ*_ ≈ 1). Conversely, when direct selection dominates background selection (|*γ*| ≫ 2*Ns*_*b*_; Figure 1C), the focal allele is purged by its own fitness cost before the background can be removed. In this limit, BGS is irrelevant, and the heterozygosity is naturally unaffected. We can therefore rely on this reduced local *N*_*e*_ approximation to model the underlying allelic dynamics across the full range of selection coefficients.

**Figure 1.**
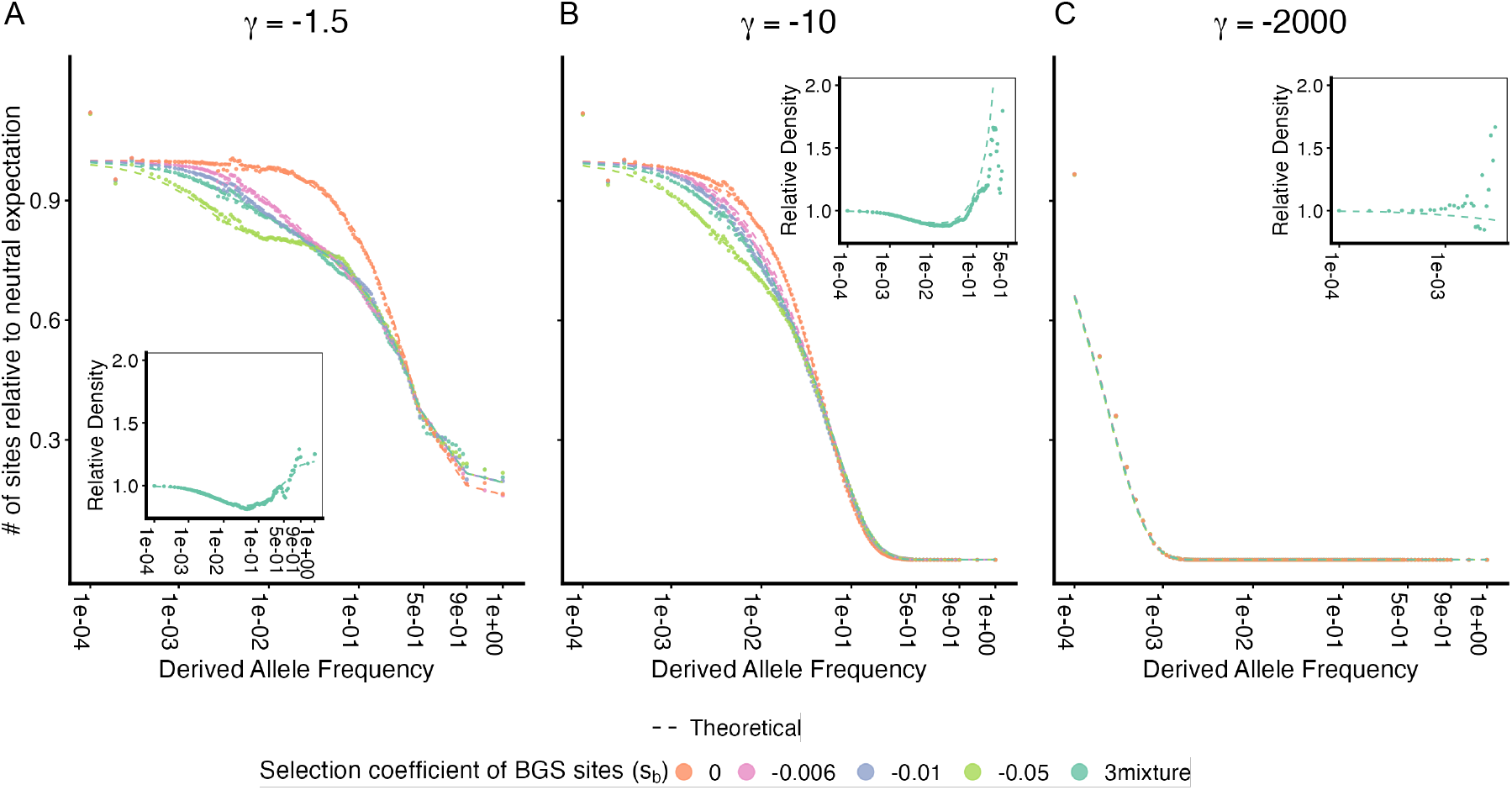
The frequency spectrum of focal selected alleles in the weak mutation limit (*λ*≪ 1). The site frequency spectrum observed in simulations (points) are compared to theoretical predictions (lines) for focal alleles under varying strengths of direct selection, assuming a background selection intensity of *B*≈ 0.82 (the human average). The site frequency spectrum is scaled relative to the standard neutral expectation (*θ/q*). Focal alleles are (A) nearly neutral (2*Ns* = −1.5), (B) moderately deleterious (2*Ns* = −10), and (C) strongly deleterious (2*Ns* = −2000). Different colors represent simulations where the same total reduction in effective population size (*B*) is achieved using different selection coefficients or a distribution with an equal mixture of all three for the background deleterious mutations (*s*_*b*_). The dashed lines represent the approximation developed in Equation 4 for panels A and B, and Equation (5) for panel C. Insets show the relative density of alleles across frequency bins, comparing scenarios with background selection to those without, with theoretical prediction plotted in dashed line.

#### Phenotype model and chromosome structure

To apply these insights in the context of phenotypic evolution, we assume a simple additive model where each site *ℓ* has two alleles differing by effect *a*_*ℓ*_. Individual *i*’s trait value is

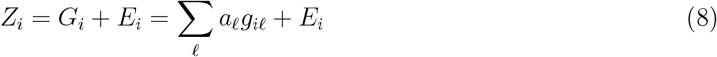

where *g*_*iℓ*_ ∈ {0, 1, 2} is the genotype count and *E*_*i*_ ∼ *N* (0, *V*_*E*_). We assume that causal sites are sufficiently spaced such that their local evolutionary dynamics can be treated independently. Each site is embedded in a recombining background that generates unconditionally deleterious alleles according to the model described above; however, we allow the density of these mutations per unit recombination (i.e. *λ* = 2^*ν*^*/*_*r*_)—and thus the magnitude of the local effective population size reduction (i.e. *B* = *e*^−*λ*^)—to vary across causal sites.

#### Exponential selection

The simplest model of directional selection is one in which fitness declines exponentially with increasing phenotype, i.e., 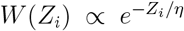 (Figure 2B). In this model, sites evolve independently with an approximately constant directional selection coefficient *s* ≈ *aη*^−1^, corresponding to a scaled selection coefficient *γ* = 2*Ns* ≈ 2*Naη*^−1^ (see Supplementary Text B).

At mutation-selection-drift equilibrium, fixations of trait-increasing and trait-decreasing mutations reach a detailed balance (Li 1987; Sella and Hirsh 2005; Rice et al. 2015). Among sites with scaled selection coefficient *γ*, the fraction of sites fixed for the trait-increasing (deleterious) allele is determined by the ratio of fixation rates,

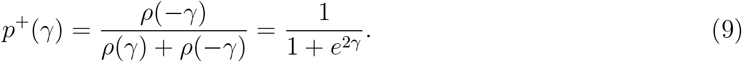

The resulting genetic variance refelcts a combination of contributions from alleles under positive and negative selection

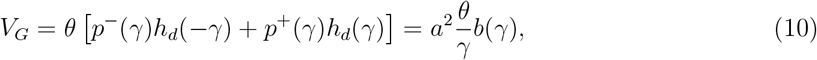

where *b*(*γ*) = (*ρ*(*γ*)−*ρ*(−*γ*)) */*(*ρ*(*γ*)+*ρ*(−*γ*)) = tanh(*γ*) measures the asymmetry of the mutational input at equilibrium.

Because of the independence among sites, the impact of background selection on each site is captured entirely by the local reduction in effective population size. This leads to a rescaled fixation asymmetry *p*^+^(*γB*) = (1 + *e*^2*γB*^)^−1^, which drives a large shift in the mean phenotype. For weakly selected sites, this shift is substantially larger than the scale of the standing genetic variation (i.e., 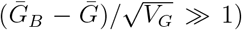, resulting in a significant decline in the mean fitness of the population (see Supplementary Text B).

**Figure 2.**
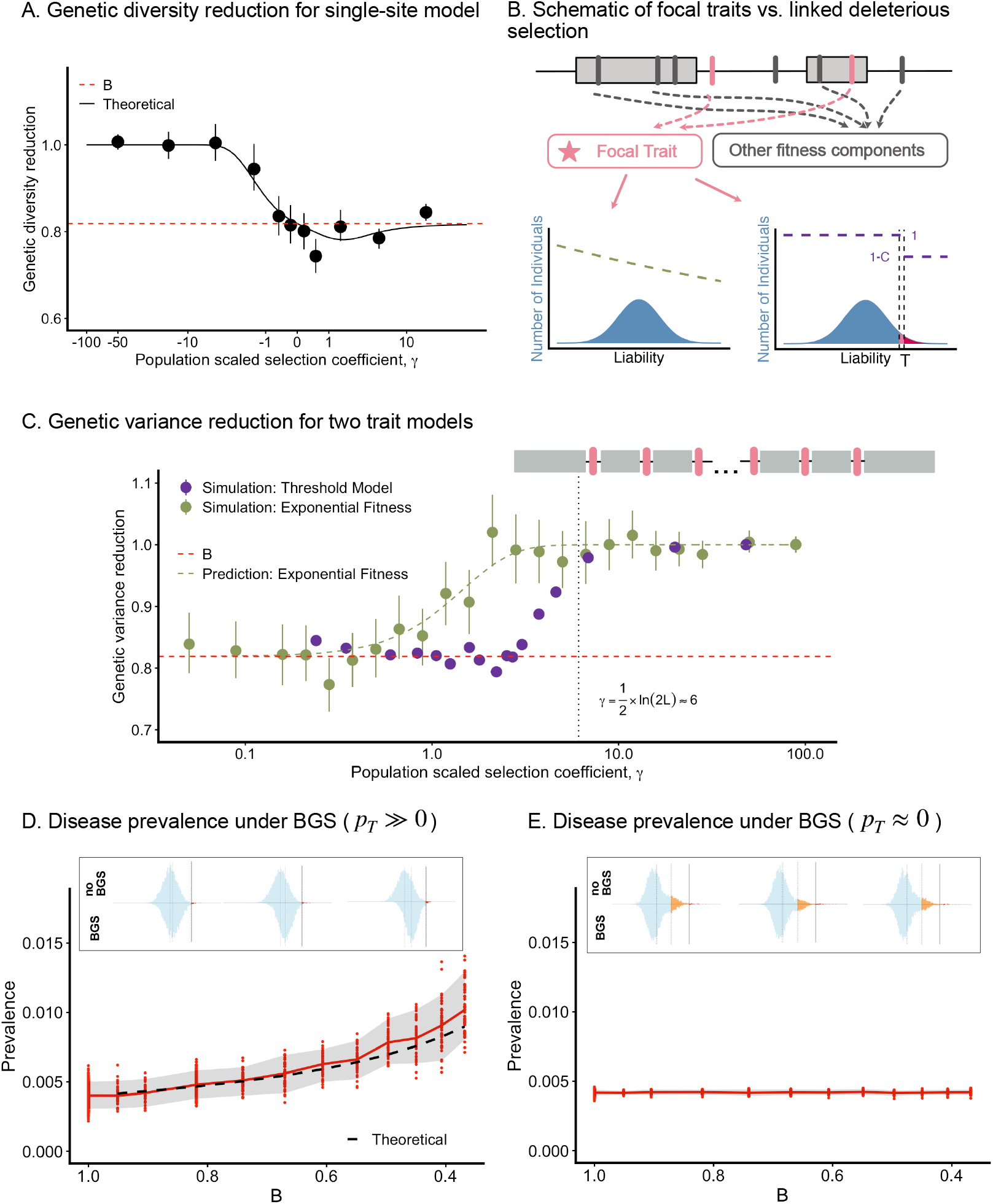
The impact of background selection on genetic variation under directional selection. (A) The reduction in genetic diversity for single-site additive directional selection model, represented as the ratio of expected heterozygosity with BGS to that without BGS (*h*_*d*_(*γB*)*/h*_*d*_(*γ*)), is plotted against the population-scaled selection coefficient (*γ*). Simulation results (points) are shown alongside the theoretical prediction from equation (7) (solid line). The horizontal red dashed line indicates the expected diversity reduction for a neutral allele, *B*≈ 0.82. (B) A schematic depicting the relationship between sites that are causal to a focal trait (pink bars) and linked deleterious mutations that contribute to other fitness components (gray bars), which are responsible for BGS effects. (C) We compare the genetic variance reduction (*V*_*G,B*_*/V*_*G*_) for the exponential fitness model (green) and the single effect size liability threshold model (purple). Simulation results are plotted as points, and the prediction for the exponential fitness function from equation 11 is plotted as a green dashed line. The horizontal red dashed line indicates the expected diversity reduction for a neutral allele, *B* ≈ 0.82. The vertical dotted line marks *γ*≈ 6, which corresponds roughly to *p*_*T*_≈ 1*/*2*L* given that we use *L* = 10^5^ to simulate. The chromosome structure we used to simulate the threshold model is depicted at right top, showing trait loci (pink bars) interspersed among BGS sites (gray regions). (D-E) Disease prevalence under BGS is shown for scenarios where (D) the equilibrium fraction of liability increasing fixations is large (*p*_*T*_ ≫ 0), versus (E) scenarios where liability increasing fixations are absent (*p*_*T≈*_ 0). In (D), the prediction from equation 17 is plotted in dashed line. In the inset, the liability distribution in one simulation is plotted comparing BGS to no BGS at background selection intensities of approximately *B* ≈ (0.95, 0.67, 0.45) from left to right.

The impact on the variance follows directly from the relationship between heterozygosity and fixation rates established in Eq. (7). Specifically, the variance is reduced by a factor

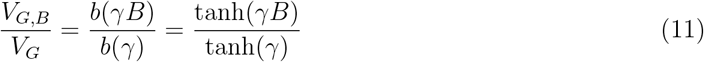

(Figure 2C; McVean and Charlesworth 1999). This reduction is greatest for effectively neutral alleles (*V*_*G,B*_*/V*_*G*_ ≈ *B* for *γ* ≪ 1) but weakens as selection strength increases, vanishing entirely for strongly selected sites (*γ* ≫ 1). This fading reflects the fact that strong negative selection purges alleles faster than BGS can remove the background, rendering the reduction in *N*_*e*_ irrelevant. Critically, because the selection coefficient *s* in this model is fixed by the fitness function parameter *η*, it is unaffected by the shift in the mean phenotype. As a result, all sites respond to their local BGS effects independently, and the effect on the phenotype in any given case is simply found by averaging these effects over the joint distribution of effect sizes and local effective size reductions.

#### Threshold epistasis

In this model, fitness depends on whether the phenotype exceeds a threshold, *T* . Individual *i*’s relative fitness is *W* (*Z*_*i*_) ∝ 1 – 𝟙[*Z*_*i*_ *> T* ]*C* where *C* is the cost of exceeding the threshold and 𝟙 [*Z*_*i*_ *> T* ] indicates whether *Z*_*i*_ *> T* (Figure 2B). We treat this as a model of complex disease, where *Z*_*i*_ represents an individual’s liability (Berg et al. 2025). Equivalent threshold models have appeared in other contexts, where the phenotype is simply termed “fitness potential” (Milkman 1978; Kondrashov 2018). We adopt the complex disease framework here, though our results could easily be applied in other contexts by relabeling terms.

##### A single effect size and a single B value

The threshold model differs from the exponential model in two fundamental ways, which are easiest to see if we first assume a single effect size, *a*, and constant local reduction in effective population size, *B*, across sites. First, the threshold imposes strong epistasis: a site’s selection coefficient depends on the variation at all other sites. However, because the background phenotypic variation smoothes out the sharp epistasis of the threshold, the marginal effect of an allele on fitness is nonetheless constant, allowing individual sites to evolve as if under additive selection. If the liability effect *a* is small relative to the variance in liability 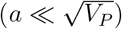, this dependency is mediated solely by the liability density at the threshold, *f* (*T*):

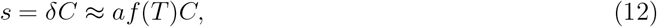

where *δ* ≈ *af* (*T*) is the site’s average effect on disease risk (Berg et al. 2025).

Second, because the threshold has a fixed position along the liability continuum, it determines the fraction of sites fixed for the liability-increasing allele 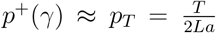 . Consequently, the threshold position determines the scaled selection coefficient via the relationship between the fixation asymmetry, i.e., *p*^+^(*γ*), the fixation rates *ρ* (*γ*) and *γ* (Eq. (9)). In a single-effect model, we can solve directly for:

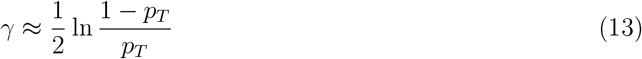

(Berg et al. 2025). Combining this with the relationship between threshold density *f* (*T*) and the selection coefficient (Eq. (12)), equilibrium is established when the mean liability is positioned at a distance just below the threshold, such that 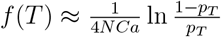 . Because sites evolve under additive selection, genetic variance depends on *γ* exactly as in the exponential model (Eq. (10)); however, *γ* is now determined by the threshold position rather than the rate of fitness decline.

In the threshold model, strong selection against individuals with liability greater than *T* prevents the sort of large shift in the mean that occurs in the exponential model. Consequently, the system must maintain the equilibrium fraction of liability increasing fixations (*p*^+^(*γ*) ≈ *p*_*T*_). As a result, BGS must reduce the selective fixation rates for liability increasing and liability decreasing mutations by the same fraction. This constraint imposes a specific requirement on the dynamics: the reduction in genetic variance must arise *solely* from the initial culling of new mutations (i.e., the factor *B*). Recall from Equation (7) that the total impact of BGS is the product of this initial cull and the change in the per-mutation contribution to variance (*h*_*d*_(*γ*_*B*_)*/h*_*d*_(*γ*)). Therefore, for the system to maintain equilibrium via the initial cull alone, the per-mutation contribution must remain unchanged. This implies that the scaled selection coefficient itself must be invariant, i.e. *γ*_*B*_ = *γ*, which then satisfies the initial requirement that the selective fixation rates are modified by the same factor, i.e.

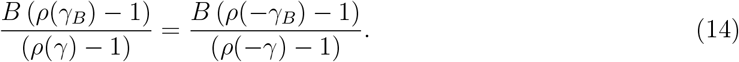

The system satisfies this requirement through a global compensation mechanism. In response to the local increase in genetic drift, the mean liability shifts closer to the threshold by a tiny amount, increasing the threshold density to *f*_*B*_(*T*) ≈ *f* (*T*)*/B*. This global adjustment increases the unscaled selection coefficients (*s*_*B*_ ≈ *s/B*), which exactly offsets the local reduction in effective population size (*N* → *NB*):

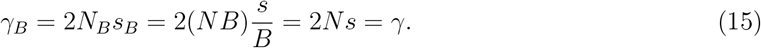

Because *γ* is invariant, the per-mutation variance *h*_*d*_(*γ*) remains unaffected, as predicted. The total reduction in genetic variance is therefore driven entirely by the reduced supply of mutations (*θ* → *θB*), resulting in a simple, *γ*-independent reduction:

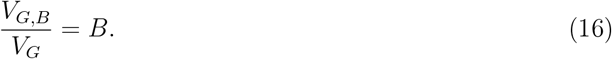

Thus, the threshold model avoids the large decrease in mean fitness characteristic of the exponential model through a global compensation of selection coefficients that maintains the mean phenotype. This compensation impacts the liability distribution by simultaneously reducing the variance (*V*_*G*_ → *BV*_*G*_) and increasing the threshold density (*f* (*T*) → *f* (*T*)*/B*) (Figure 2C, Figure S3). These two effects combine to increase the disease prevalence (Figure 2D) by a factor of approximately

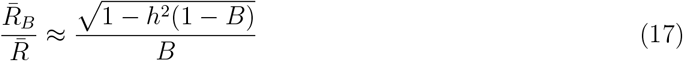

where *h*^2^ is the heritability of liability in the absence of BGS (We also use simulations to explore whether the skew in the frequency spectrum impacts the prevalence, but observed no effect; see Supplementary Text C, Figure S12).

However, both the invariance of *γ* and the increase in prevalence depend entirely on the global compensation mechanism, which is itself predicated on the existence of deleterious fixations to balance (i.e., *p*_*T*_ *>* 0; Eq. (13)). This assumption breaks when the threshold is so low that *p*_*T*_ is driven to effectively zero. In a genome with *L* contributing sites, this occurs roughly when the expected number of fixed deleterious alleles drops below one (*p*_*T*_ ≲ 1*/*2*L*), corresponding to a critical selection coefficient of 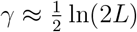. If *γ* exceeds this value, there are no deleterious fixations left to maintain. The global compensation mechanism shuts down, *γ* is no longer forced to be invariant, and the system reverts to standard local dynamics. Consequently, in this strong selection regime, the distinctive signatures of the threshold model disappear: the *γ*-independent variance reduction fades away (Figure 2C), and the increase in disease prevalence predicted by Equation (17) does not occur (Figure 2E).

##### Variation in effects and variation in B values

The threshold model described above assumes a single effect size and constant *N*_*e*_ reduction, allowing for a straightforward population size rescaling. However, if effect sizes (*a*) and local *N*_*e*_ reductions (*B*) vary across sites, the dynamics are complicated by the fact that all sites remain coupled by a single threshold density, *f* (*T*). The BGS-driven increase in *f* (*T*) must therefore represent a global compromise set by the varying impacts of local effective size reductions across sites. This single compensatory change couples back to all sites, impacting their individual selection coefficients. To capture these dynamics, we must first determine how the global compensatory evolution of *f* (*T*) is determined, and then resolve how this common shift interacts with local effective size reductions across the distribution of effect sizes.

For simplicity, suppose *a* and *B* vary independently across sites with densities *g*_*a*_ (*a*) and *g*_*B*_ (*B*). Then, the equilibrium threshold density, *f* (*T*), must solve:

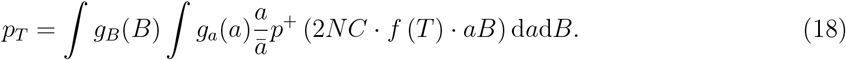

In Supplement F, we find analytical approximations for this integral in tractable limits assuming a gamma distribution on effect sizes *g*_*a*_(*a*). Here, we summarize the resulting behaviors by categorizing the distinct regimes controlled by the value of *p*_*T*_ (Figure 3B). We first consider the effectively neutral limit 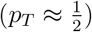 before examining the transition toward the strong selection limit (*p*_*T*_ → 0). Crucially, as selection strengthens, the solution depends heavily on the shape of the effect size distribution; we therefore contrast distributions with a low coefficient of variation (CV) against those with a high CV (e.g., “L-shaped” distributions) to illustrate how the tail of the distribution shapes the global compensation.

**Figure 3.**
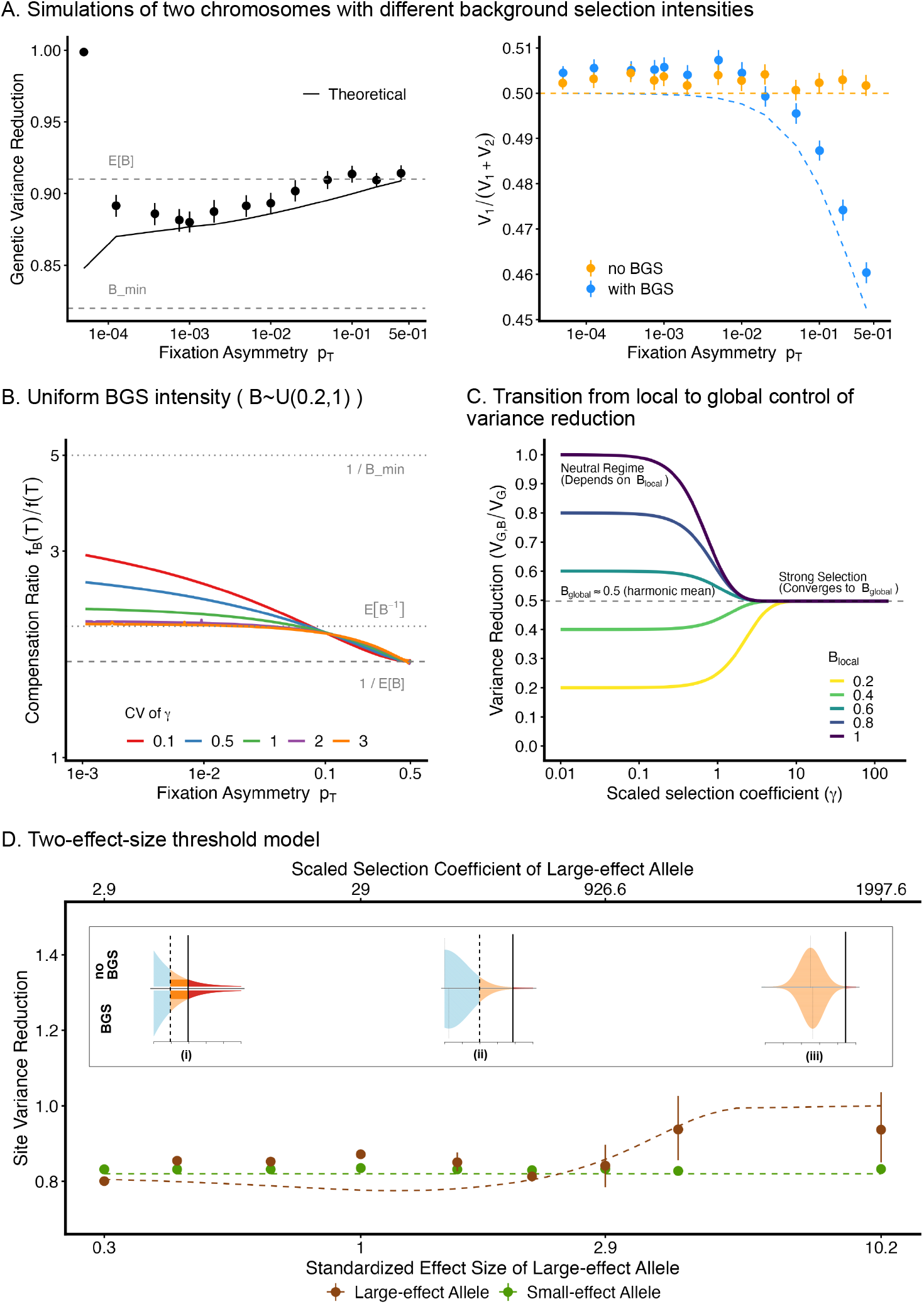
Global coupling of background selection effects in the liability threshold model. (A) We simulated using a single effect size model with causal sites evenly distributed across two chromosomes, and varied the fixation asymmetry by varying the relative position of the threshold, *p*_*T*_ . In one set of simulations, there is no BGS. In the second set of simulations, one chromosome experiences an effective size reduction due to BGS of *B ≈* 0.82 while the other experiences no BGS. The left panel reports the total reduction in genetic variance (across both chromosomes)in the simulations with BGS versus those without. The solid black line plots the numerical solution of equation 18. The right panel reports the fraction of genetic variance contribute by chromosome one in the presence (blue) and absence of BGS (orange). The theoretical prediction is obtained from equation 22. (B) The ratio of threshold density, *f*_*B*_(*T*)*/f* (*T*), is plotted against the fixation asymmetry (*p*_*T*_) for different coefficients of variation (CV) of the effect size distribution. Curves represent numerical solutions to equation 18, assuming a Gamma distribution for effect sizes and a uniform distribution of local B values, *B*∼ *U* (0.2, 1). Horizontal dashed lines indicate theoretical limits derived in the text. (C) The variance reduction due to BGS as a function of scaled selection coefficient *γ* and local *B* value (i.e. Eq. (22)), assuming *B* ∼ *U* (0.2, 1) and the high CV limit such that *B*_*global*_ = 1*/*𝔼 [*B*^−1^]. Curves illustrate the transition from dependence on *B*_*local*_ to the global average governed by *B*_*global*_. (D) We simulated a two effect size architecture where one set of sites has a small effect size and belongs to the the weak selection regime, while the other has a large effect size and belongs to the strong selection regime (all sites have the same local effective size reduction). We varied the effect sizes of the large-effect alleles. The x axis on the bottom of the figure measures the size of the large effect relative to the phenotypic standard deviation, while the axis on the top of the figure measures it in terms of the population scaled selection coefficient, *γ*. The pattern of genetic variance reduction for all but the largest effect sites illustrates the spreading of BGS effects to strongly selected sites via the global compensation mechanism. The theoretical predictions are obtained by solving the two effect model numerically, see Supplementary section D (i) – (iii) illustrate how the risk effect size, *δ*, for the large-effect-size allele changes with BGS depending on its effect size and drive this effect (recall that *s ≈ δC*). The region illustrates the proportion of the population who are pushed across the threshold by the allele, and thus its effect on disease risk and fitness. (i) The effect size is relatively small and the risk effect size can be linearly approximated via the dark orange rectangle. BGS increases the area of this rectangle by a factor of 1*/B*. (ii) This linearity breaks down, and the effects on risk and therefore fitness increase by more than 1*/B*. (iii) When the effect size is large enough to push the entire population across the threshold, BGS no longer has any impact.

When 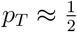, neutral dynamics and local *N*_*e*_ effects dominate. The global compensation required to maintain equilibrium is set by the arithmetic mean of *B*:

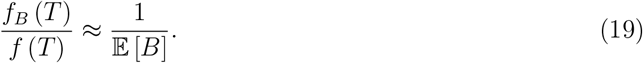

However, because most sites are effectively neutral, they are insensitive to this global change in *f* (*T*). Their dynamics are instead dominated by their local *N*_*e*_ reduction, meaning their individual variance reductions simply depend on their local *B* value.

Now consider the limit where we decrease *p*_*T*_ toward zero. The solution to Equation (18) depends heavily on the shape of the effect size distribution, *g*_*a*_(*a*). Suppose *g*_*a*_(*a*) has a low coefficient of variation (CV), meaning effect sizes are tightly clustered around the mean. As *p*_*T*_ decreases, the bulk of the sites transition into the strong selection regime simultaneously. In this limit, the fixation asymmetry decays exponentially (i.e., *p*^+^(*γ*) ≈ *e*^−2*γ*^). The fixed liability burden at equilibrium is therefore sustained almost entirely by the few sites where genetic drift is strong enough to overpower selection—specifically, those with the smallest scaled selection coefficients. In a model where effect sizes are all roughly equal to one another, these sites are the ones with the strongest local BGS effects (i.e. the smallest *B* values). Consequently, the global compensation converges to depend on the minimum *B* value in the genome:

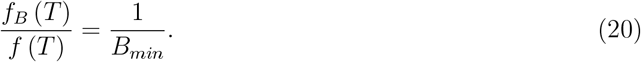

Thus, in this limit, the specific local *N*_*e*_ reductions at most sites become irrelevant, as the system’s global equilibrium is determined by the region experiencing the strongest background selection.

However, as in the single-effect-size case, this compensation mechanism shuts down when 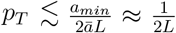 . This occurs because at this point, even sites with the minimum effect size, *a*_*min*_, can no longer be fixed for the deleterious allele. Thus, for a fixed low-CV distribution, decreasing *p*_*T*_ from 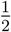 to 0 causes the ratio *f*_*B*_ (*T*)*/f* (*T*) to first increase from 𝔼 [*B*]^−1^ toward 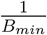, and then abruptly drop to 1 as *p*_*T*_ approaches 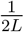 (Figure 3A).

Alternatively, if the effect size distribution has a high coefficient of variation (CV)—corresponding to the “L-shaped” limit of the gamma distribution—sites will be spread across all selective regimes simultaneously. While the long tail extends into the strong selection regime, the bulk of sites remain in the effectively neutral regime near zero. The resulting global compensation factor, *f*_*B*_ (*T*)*/f* (*T*), must therefore represent a compromise between the limits set by these different regimes: the large mass of neutral sites pulls the compensation toward the neutral limit (𝔼 [*B*]^−1^), while the exponential sensitivity of the sites in the tail pulls it toward the strong selection limit 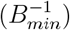. In Supplement F, we show that in this high-CV limit, this compromise converges to the reciprocal of the harmonic mean *B* value:

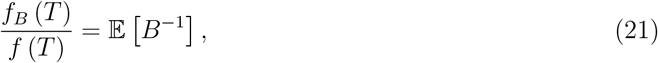

which satisfies 1*/*𝔼 [*B*] ≤ 𝔼 [*B*^−1^] ≤ 1*/B*_*min*_ for any distribution *g*_*B*_. In Figure 3B, we illustrate the convergence to this limit by solving equation (18) numerically assuming the local effective size reductions experienced by causal sites are uniformly distributed between 0.2 and 1.

In contrast to the low-variance case, this global compensation persists even as *p*_*T*_ → 0. This difference stems from the high density of small-effect variants in distributions with a mode at zero (e.g., the high-CV Gamma). In the low-CV case, all sites have approximately the same effect (*a* ≈ *ā*), so they all transition into the strong selection regime simultaneously, causing the compensation mechanism to shut down abruptly. In the high-variance case, there is no such simultaneous shutdown. As the threshold density *f* (*T*) increases, sites with large effects are indeed pushed into the strong selection regime, where they stop contributing to the fixed liability burden. However, the mode at zero ensures a continuous reservoir of sites with even smaller effects. As sites with larger effects are all pushed into the strong selection regime, the burden of maintaining the total equilibrium fixation asymmetry dictated by the threshold shifts to these smaller effect variants, which—precisely because of their small size—remain in the weakly selected regime where they are sensitive to changes in local effect size. Thus, baring pathological cases (see Supplementary Text F), this supply of weakly selectedsites is effectively inexhaustible, maintaining the global compensation factor at *f*_*B*_ (*T*)*/f* (*T*) ≈ 𝔼 [*B*^−1^] as *p*_*T*_ → 0.

While the specific result in Equation (21) arises from the power-law tail of the high-CV gamma distribution, the broader insight is that in biologically realistic scenarios—where both effect sizes (*a*) and local *B* values vary—the phenotype is shaped by a combination of two forces: a single global compensation factor and site-specific local BGS effects. The total impact on any given site depends on its selective regime. For a site with scaled selection coefficient *γ* and local *N*_*e*_ reduction *B*_*local*_, the total reduction in genetic variance is given by:

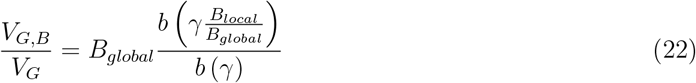

where *b*(*γ*) = tanh(*γ*) is the mutational asymmetry and *B*_*global*_ = *f* (*T*)*/f*_*B*_(*T*) is the global compensation factor determined by the genome-wide distributions of effects and *B* values (e.g., in the high-CV gamma limit, *B*_*global*_ is the harmonic mean, 1*/*𝔼 [*B*^−1^]). This implies that the system behaves in two distinct ways depending on the selection regime: effectively neutral sites are governed by their local effective size reductions (*V*_*G,B*_*/V*_*G*_ ≈ *B*_*local*_), while strongly selected sites are pinned to the global average (*V*_*G,B*_*/V*_*G*_ ≈ *B*_*global*_), regardless of the local *B* value. We illustrate this transition in Figure 3C for the same uniform distribution of local effective size reductions used in Figure 3B.

Notably, this result demonstrates that BGS reduces genetic variance even at sites under very strong selection. In the single-effect model, such sites were immune to BGS because the global compensation mechanism shut down entirely at the *p*_*T*_ → 1*/*2*L* boundary. By contrast, in a model with a high-variance distribution of effect sizes, the global compensation persists, and effectively subjects these strongly selected sites to a variance reduction dictated by the genome-wide background. This globally driven effect, however, has its own limit. The underlying approximation *s* ≈ *af* (*T*)*C* eventually breaks down for sites with very large effects, as their selection coefficients begin to scale non-linearly with their liability contributions (Berg et al. 2025). For sites with effects comparable to the standard deviation of liability 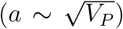, this non-linearity can even drive variance reductions that are greater than *B*_*global*_, as the selection coefficients increase faster than linearly with increasing threshold density in this range. As *a* increases further, however, the coupling breaks down entirely: sites with effects so large that carriers cross the threshold regardless of their genetic background have selection coefficients determined solely by the fitness cost (*s* ≈ *C*), independent of the threshold density *f* (*T*). Thus, for sites under the very strongest selection, the impact of BGS does ultimately fade away. Crucially, the point at which this occurs is not determined by population-genetic limits on fixation dynamics, but by the fitness cost of the threshold itself. We illustrate these effects in Figure 3D by comparing numerical solutions (see Supplementary Text D) with simulations of models with two effect sizes, varying the magnitude of the larger effect relative to the background variance in liability.

This result demonstrates that in a realistic model with a high-variance distribution of effect sizes, BGS reduces genetic variance even at sites under very strong selection. Because the wide distribution of effects ensures that global compensation (*B*_*global*_) is active, even strongly selected sites remain coupled to the genome-wide background and experience a variance reduction dictated by it. This stands in contrast to the single-effect model, where such sites are immune to BGS because the entire compensation mechanism shuts down abruptly once the unique effect size transitions into the strong selection regime (at the *p*_*T*_ → 1*/*2*L* boundary).

This globally driven effect, however, has its own limit. The underlying approximation *s* ≈ *af* (*T*)*C* eventually breaks down for sites with very large effects, as their selection coefficients begin to scale non-linearly with their liability contributions (Berg et al. 2025). For sites with effects comparable to the standard deviation of liability 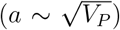, this non-linearity can drive variance reductions even greater than *B*_*global*_. As *a* increases further, however, the coupling breaks down entirely: sites with effects so large that carriers cross the threshold regardless of their genetic background have selection coefficients determined solely by the fitness cost (*s* ≈ *C*), independent of the threshold density *f* (*T*). Thus, for sites under the very strongest selection, the impact of BGS does ultimately fade away.

However, the point at which this occurs is not determined by population-genetic limits on fixation dynamics, but by the fitness cost of the threshold itself.

### Stabilizing selection

As we have shown, fitness epistasis in the form of threshold selection alters the impact of BGS via a global compensation mechanism, which is necessary to maintain the specific fixation asymmetry imposed by the threshold. We now turn to a model of stabilizing selection, defined by a Gaussian fitness function with an optimum phenotype of *Lā* (i.e., 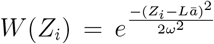 ; Lande 1975). This model contrasts with the threshold case in two key ways. First, it ensures mutational symmetry at equilibrium, eliminating the net fixation asymmetry that drove the global coupling in the threshold model. Second, the local dynamics differ: this form of epistasis causes individual alleles to evolve as if underdominant, with a selection coefficient 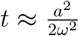 against heterozygotes (Fisher 1930; Robertson 1956). This allows us to isolate the impact of BGS on these purely local dynamics.

Defining the scaled coefficient of underdominance as *τ* = 2*Nt*, Hayward and Sella 2022 showed that the expected lifetime contribution to heterozygosity is

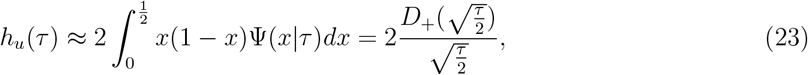

where *D*_+_(*y*) is the Dawson function, and 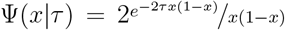 is the sojourn time at minor allele frequency *x*. The behavior of this function reveals a crucial deviation from simple directional selection. While *h*_*u*_(*τ*) converges to the directional result *h*_*d*_(−*τ*) at the extremes (*τ* ≪ 1 and *τ* ≫ 1), it is significantly elevated in the intermediate regime (1 ≲ *τ* ≲ 30). In this intermediate regime, alleles that drift to intermediate frequency “slow down” and persist longer than predicted by negative selection alone, causing the ratio *h*_*u*_(*τ*)*/h*_*d*_(−*τ*) to peak at ≈ 1.29 when *τ*^∗^ ≈ 4.42 (Figures S7 and S8).

Applying the reduced effective size approximation (*N*_*e*_ → *N*_*e*_*B*), the modification to heterozygosity is given by

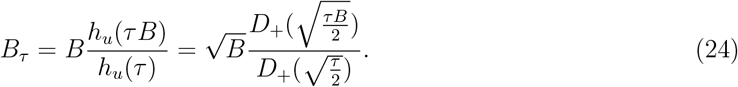

This equation demonstrates that BGS does not simply reduce variance by a factor of *B* or cancel out as it does under strong directional selection. Rather, for some combinations of *τ* and *B*, the increased rate of drift leads sites that would otherwise be held at low frequencies to drift up to higher frequencies where the selection against them is weaker, allowing them to continue drifting even further. The increased contribution to heterozygosity conditional on surviving the low frequency cull due to BGS therefore overcompensates for heterozygosity lost that is eliminated by it.

This variance inflation is maximized when BGS shifts alleles from the strong selection regime down to the intermediate regime where the impact of underdominance is most pronounced (*τ*^∗^ ≈ 4.42). For strong BGS (*B* ≪ 1), this peak net increase occurs at 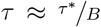 and converges to the theoretical maximum of ≈ 1.29, while the range of affected sites scales with 1*/B*. Figure 4A shows that multilocus simulations of stabilizing selectino faithfully replicate these single-site predictions.

In Figures 4C and 4D, we apply the frequency dependent factor *B* (*q*) from equation (2) and the rescaling *τ* → *τ B* to the frequency spectrum for underdominant sites for *τ* = −1.5 and 10 under modest background selection (i.e., *B* = 0.83, the human average). In this regime, the rescaling of *τ* is not dramatic enough to move sites from the strong into the weak selection regime, so the effects on the frequency spectrum are modest. In contrast, under larger effective population size reductions, as BGS moves sites fully between regimes, the inflation of the frequency spectrum at intermediate frequencies becomes substantial, although here our simple approximation for the skew in the low frequency regime breaks down (Figure S9).

## Discussion

In this paper, we have used a combination of analytical theory and simulations to study the impacts of background selection in recombining genomes, both at the level of individual variants and the level of phenotypes, in several long-studied models.

### Background selection and the frequency spectrum in recombining genomes

While the impact of background selection on aggregate diversity is often well-approximated by a simple reduction in effective population size, characterizing the full shape of the site frequency spectrum in recombining genomes presents a more difficult challenge. Here, we describe an effectively nonrecombining block approximation (Neher et al. 2013; Good et al. 2014; Weissman and Hallatschek 2014) for the classic model of background selection with recombination by defining the characteristic length scale of the process, and demonstrate that this approximation recovers classic results previously derived by summing contributions across many loci in pairwise models (Hudson and Kaplan 1995; Nordborg et al. 1996). This re-framing allows us to make progress in understanding the impact of BGS on the skew in the frequency spectrum in recombining genomes and to derive a simple expression for the effect that is accurate in the weak mutation regime which characterizes humans (*λ* ≲ 1*/*2).

**Figure 4.**
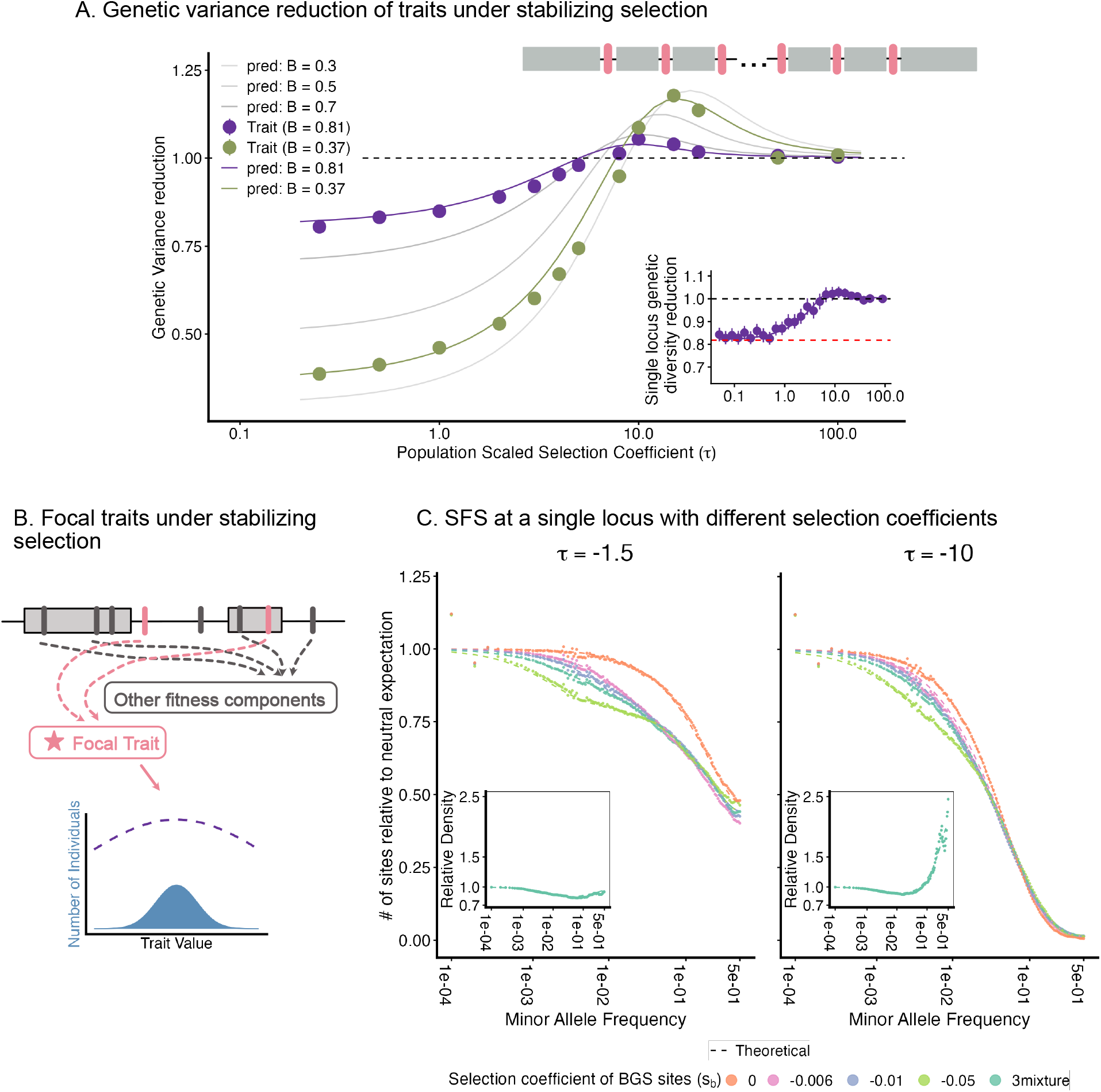
Stabilizing Selection. (A) We simulate traits with a single effect size under stabilizing selection with two different effective population size reductions. For each effective size reduction, we plot the change in genetic variance against the population-scaled selection coefficient 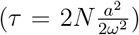. The chromosome structure used for simulation is depicted in the top right, showing trait loci interspersed among BGS regions. Simulations results (points) are compared with the theoretical prediction, obtained from equation 23, with the B values estimated from the diversity reduction of neutral sites simulated in the same genome. The inset shows the genetic diversity of underdominant alleles in a single locus simulations with *B* = 0.82, the human average. (B) Schematic describing the relationship between sites contributing to focal traits (pink bars) and linked deleterious mutations (gray bars). (C) The site frequency spectrum observed in simulation (points) are compared to the theoretical predictions (dashed lines) for focal selected sites under weak underdominant selection (*τ* = − 1.5, left) and stronger underdominant selection (*τ* = − 10, right). Different colors represent different selection coefficients or a distribution with an equal mixture of all three at the background mutations (*s*_*b*_). Insets show the relative density of alleles at each frequency bin with background selection vs. no background selection.

In Supplementary Text A, we use heuristic arguments to extend the effectively non-recombining block approximation into a “shrinking block” model. In this framework, the length of the original haplotype remaining associated with a focal allele of age *t* is *M*_*t*_ ≈ 2*/rt*. This approximation allows us to map the dynamics onto the non-recombining model of Cvijović et al. (2018) and extend their piecewise expression for the frequency spectrum skew under *λ* ≫ 1 to recombining genomes (Figure S5).

Notably, this shrinking block perspective reveals a key difference in dynamics between the two models: because *M*_*t*_ is large for small *t*, an allele in a recombining genome becomes associated with newly arising deleterious mutations far beyond the length 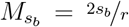 that characterizes classical background selection (BGS) against pre-existing variation. Over the short timescales that characterize this process, the balance between mutation and recombination ensures that the number of new deleterious mutations associated with the focal allele remains constant on expectation and is also equal to *λ* (Eq. (A.9)). Consequently, the time required for an allele to feel the effect of selection against new mutations on its background is reduced by a factor of 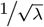 relative to a non-recombining model with the same local reduction in effective size (Eq. (A.17)). This implies that the frequency at which substantial skew begins to accumulate is lowered by the same factor (Eq. (A.18)), and that the blocks eliminated by this process are longer than the characteristic length 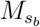 by a factor of *λ* (Eq. (A.12)).

Developing a complete analytical description of the frequency spectrum for *λ* ≳ 1*/*2 (or a numerical method to compute it accurately) remains a priority for future work. For 1*/*2 ≲ *λ* ≲ 1, progress might be made by assuming the focal allele accumulates at most one additional deleterious mutation within the characteristic length 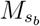, though this approximation would clearly fail for larger *λ*. For the *λ* ≫ 1 case, the shrinking block perspective is promising: the impact on an allele at frequency *q* could be described by a characteristic length *M*_*q*_ = (*Nrq*)^−1^, treating the region as a non-recombining chromosome of that length. While this approach requires a more robust solution for the frequency spectrum in non-recombining genomes, we hope these insights can help provide a bridge toward that solution.

### Models of phenotypic selection and linkage in recombining genomes

In our phenotypic analyses, we assume 2*NBs*_*b*_ ≫ 1, ensuring that the skew in the frequency spectrum is fully “priced in” at relatively low frequencies. This permits us to assume that BGS preserves the standard functional forms of heterozygosity and fixation rates under Wright-Fisher diffusion, allowing us to apply classic effective population size rescaling in our phenotypic models while retaining the standard . However, the relationship between an allele’s contribution to heterozygosity and its fixation rate expressed in equation (6) is more general than the diffusion limit, as it simply represents an averaging of Fisher’s fundamental theorem over an allele’s transit through the population (Fisher 1930; Santiago and Caballero 2016). Consequently, the behaviors we describe for directional models are expected to be extremely general, and should equally apply in models that relax the strong selection assumption (Good et al. 2014; Santiago and Caballero 2016; Barroso and Ragsdale 2025). Under the threshold model in particular, introducing essentially any form of linked selection should force the population distribution to evolve such that the selective fixation rates in both directions are reduced by the same factor. Even if the form of linked selection alters the functional form of the fixation rate, a reduction in the selective fixation rate by a given multiplicative factor must imply a corresponding reduction in heterozygosity by the same factor, independent of the underlying details (although changes in the form of the fixation rate could alter the precise form of *B* value averaging that drives the global effect).

That said, different models of linked selection nonetheless result in different relationships between fundamental parameters and patterns of genetic diversity, and there remains considerable uncertainty about the underlying source of linked selection signals (Murphy et al. 2022; Buffalo and Kern 2024). While we have primarily focused on understanding the impact of BGS on the equilibrium evolution of complex traits, it is also interesting to ask the converse question: what forms of phenotypic variation underlie linked selection signals that have been measured empirically, and what model of selection applies to the underlying loci? This is particularly interesting to consider in light of arguments for the ubiquity of stabilizing selection (Simons et al. 2018; Koch et al. 2024; Simons et al. 2025).

Intuitively, alleles with effects large enough to fall into the strong selection regime (i.e., 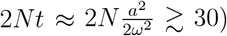 should impact linked sites precisely as predicted by classic BGS theory, because their local dynamics mirror the standard strong selection model. A more complex question is what effect we should expect from loci falling into the intermediate weak selection regime, where underdominant dynamics become important. These loci are more likely to drift to intermediate frequencies and persist than their counterparts under weak *directional* selection (which can be modeled using the quantitative genetic framework of Santiago and Caballero 2016). However, they also convert additive variance into various forms of interaction variance as they change frequency, and it is unclear to what extent this non-additive variance might remain correlated with the focal allele across generations, or how the conversion of variance between different statistical partitions impacts linked selection dynamics. This remains an interesting avenue for future work.

Of course, the linked selection effects we study are not the only mechanism through which linkage influences complex trait evolution, particularly under stabilizing selection. By selecting against extreme phenotypes, stabilizing selection generates negative LD between alleles with similar effects, causing a short-term reduction in genetic variance known as the Bulmer effect (Bulmer 1971; Bulmer 1974). From the perspective of a single focal site, the Bulmer effect manifests as a negative covariance between the trait-increasing allele and the genetic background. This attenuates the allele’s marginal phenotypic association and, by extension, its marginal fitness effect (i.e., it decreases the site’s “effective” selection coefficient; Negm and Veller 2024). Although not the primary focus of our study, we found while calibrating our simulations that this reduction in selection strength leads to a longterm increase in *genic* variance that exceeds the short-term reduction caused by negative LD. Thus, if stabilizing selection is sustained consistently over the timescales on which individual alleles make their transits through the population, then the net impact of linkage may actually be to increase the genetic variance of quantitative traits, rather than to decrease it (see Supplementary Text E), even before accounting for the potential variance increasing effects of BGS that we describe.

### Implications for natural populations

The extent to which background selection shapes genetic architecture in nature depends critically on the underlying model of phenotypic selection and/or the marginal fitness effects of individual loci. Naturally, for effectively neutral loci, the specific selection model is irrelevant, and the impact of BGS follows standard neutral expectations. Thus, the observation by Rockman et al. (2010) in *C. elegans* that loci in gene-poor, high-recombination regions explain significantly more gene expression variance implies that if BGS is responsible, the genetic variance must be effectively neutral. The most interesting potential effects are thus for strongly selected loci, where different models diverge in their predictions.

Much of the literature on complex trait evolution assumes that stabilizing selection is the norm (Kingsolver et al. 2001; Sella and Barton 2019). While this assumption has historically been difficult to assess, several recent analyses in human genetics support the hypothesis, a finding that appears to hold for diseases as well as quantitative traits (Sanjak et al. 2018; Simons et al. 2018, 2022; Koch et al. 2024; Berg et al. 2025; Ragsdale 2025). Suppose we accept the maximalist interpretation of this inference and assume that all functional loci contribute to fitness variance via their effects on traits under stabilizing selection. What, then, would be the effect of BGS on functional variation? In humans, the consequences would be straightforward and relatively minor, given the relatively modest local reductions in effective population size. However, in species experiencing stronger reductions, the impact is more significant, as we would expect the contribution to variance to increase across a wide range of sites that would otherwise by strongly selected. While the increase in heterozygosity due to BGS is capped at ≈ 1.29—and thus can never provide more than a small boost to total genetic variance—the impact on genetic architecture could be substantial, as strongly selected loci, where alleles would typically be confined to low frequencies, would instead have a significant portion of their variance contribution driven by variants drifting to intermediate frequencies. Notably, this dynamic would be expected to operate only to the extent that *τ <* 2*Ns*_*b*_, so that the effects of background selection have time to take effect.

While it seems clear that stabilizing selection plays an important role in shaping much functional variation, it is reasonable to question whether it is the whole picture. It could be that axes of phenotypic integration which experienced a sustained directional mutation-selection balance are simply more difficult to measure, and have thus evaded detection. If a substantial fraction of functional variation is under directional selection, and if most directional selection were derived from traits subject to exponential fitness functions (or some other model in which individual loci have independentfitness effects), then there is no effect on the contribution to variance from strongly selected sites, although the frequency spectra may still be shifted toward large effect loci is |*γ*| *<* 2*Ns*_*b*_, though in a less extreme way than under stabilizing selection because of the absence of the slow down dynamic at intermediate frequencies.

However, models in which selection acts independently predict unreasonably large genetic load. This occurs for the same reason that the exponential model predicts large changes in mean fitness due to BGS: if loci fix independently, the accumulation of weakly selected fixations drives mean fitness down by several orders of magnitude—a result inconsistent with the persistence of natural populations. Stabilizing selection offers one resolution to this “load paradox” (Charlesworth 2013; Barton 2017). Strong synergistic epistasis of the kind present in the liability threshold model, offers another, for precisely the reason that the threshold model differs from the exponential model in our analysis: the sharp increase in the rate of mean fitness decline with increasing phenotype. (Although we frame our analysis in terms of disease, any model exhibiting this pattern will behave similarly; Kondrashov 1995; Berg et al. 2025).

Thus, to the extent that individual variants derive their fitness effects from directional mutation-selection balance, there has been a strong theoretical expectation that synergistic epistasis must be involved. Despite this, empirical support for its importance remains mixed, for a variety of technical reasons (Mukai 1964; Mukai and Cockerham 1977; Fry 2004; Halligan and Keightley 2009; Sohail et al. 2017; Garcia and Lohmueller 2021; Sandler et al. 2021). However, efforts to detect synergistic epistasis are often underpowered because they typically test for interactions between random sets of alleles, when we might expect *a priori* that it exists only among sets of variants which integrate into the same phenotype, of which there are presumably many (Rice (1998); but see Lee (2022) for a counterexample). Consequently, it remains difficult to determine how much “evidence of absence” should be inferred from the current scarcity of empirical evidence.

Thus, while the relative importance of synergistic epistasis in evolution remains unclear, our results suggest that to the extent that it is important, then it is likely that BGS plays a substantial role in reducing contributions to genetic variance from strongly selected loci which are otherwise largely immune to its local effects. These effects would be most pronounced in species where the average genome-wide reduction in effective size is large (i.e. species with a high density of deleterious mutations per unit of recombination), or where it varies substantially across different parts of the genome, given that the magnitude of the global compensation is driven more by sites with the smallest *B* values. Notably, because this phenomenon manifests through a genome-wide change in the marginal selection coefficients themselves, it would be difficult to detect empirically, other than by demonstrating that both synergistic fitness epistasis and BGS are operating.

## Methods

### Simulation of single-site selection models

To validate the predictions on diversity reduction by BGS in our single-site models, we performed SLiM simulations (Haller and Messer 2019) on a single site under directional or underdominant selection across a range of selection coefficients. In each case, we simulated 5, 000 recombining diploid genomes, with each genome having *M* = 4 × 10^5^ background sites, and a single focal site in the middle of the chromosome. The per-site recombination rate is set to 2.5 × 10^−6^, such that the map length is equal to 1 Morgan. The mutation rate at each of the M background sites is 2.5 × 10^−7^. When we simulate without background selection, all mutations at the *M* background sites other than the focal one are neutral. When we simulate with background selection, all mutations at these M background sites have a selection coefficient *s* = −0.005. With *ν* = 2.5 × 10^−7^ and *r* = 2.5 × 10^−6^, this produces a *B* of 0.82. To vary the background selection strength, we tune the mutation rate (*ν*) with a fixed recombination rate (*r*) to produce a specified 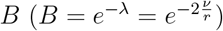.

The mutation rate at the focal site is set to zero, and we introduce mutations at this site “manually”.

We first let the simulation of the background sites run for 2*N* generations so that the population reaches mutation-selection balance. After this burn-in period, we introduced a single selected allele at the focal site in the center of the genome.

To measure the reduction of heterozygosity by BGS, we adopt the method used by Charlesworth et al. (1993) to measure the expected total contribution to heterozygosity of a new mutation during its transit through the population (*H*_*tot*_). This is given by the sum of heterozygosity at the focal site during the *T* generations that the mutation segregated in the population,

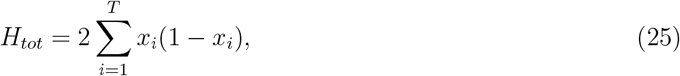

which has a theoretical expectation of 𝔼 [*H*_*tot*_|*γ*] = *h*_*d*_(*γ*), or 𝔼 [*H*_*tot*_|*τ* ] = *h*_*u*_(*τ*), depending on the model, and is thus proportional to the expected heterozygosity for a single site at a random sampling time in the low mutation limit. For each selection coefficient, we average the value of *H*_*tot*_ across 10^7^ separate focal sites. These mutations were simulated across 10^4^ replicate burn-ins, with 10^3^ mutations considered per burn-in. We then take the ratio of *H*_*tot*_ under BGS relative to no BGS as our estimator of the impact of background selection on the expected heterozygosity. To obtain a standard error, we treat the value of *H*_*tot*_ for a single mutation’s transit through the population as a single data point, and bootstrap across these values.

To construct the site frequency spectrum (SFS), we record the allele frequency of each new mutation at every generation during its sojourn in the population. To validate the frequency-dependent B approximation, we generate multiple SFS datasets in which *ν* and *r* are held constant, ensuring the same expected reduction in neutral diversity across datasets. We then vary the selection coefficients of background mutations, *s*_*b*_ ∈ {−0.006, −0.01, −0.05}, as well as an equal mixture of these three coefficients.

### Simulation of complex Traits

#### Exponential selection model

To plot the reduction in variance due to background selection under the exponential model, we take advantage of the theoretical equivalence between the exponential model and the single-site additive model. To this end, we simulate a single site under directional selection with a fixed selection coefficient, as described above, except that each time a mutation ends its transit through the population and we revert the simulation back to the end of the burn-in, we choose the sign of the effect of the next mutation to be opposite to that of the allele that fixed at the end of the previous mutation’s transit; i.e. if it was the deleterious allele that fixed, then the next mutation would be beneficial, and vice versa. This scheme simulates the long-term fixation dynamics under the exponential model. Calculations regarding the reduction in heterozygosity due to BGS are then otherwise performed as described above in the single-site case.

#### Threshold selection model

In this case, we simulate under the full threshold model, again with 5, 000 diploid individuals. For each choice of 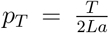 in the single-effect-size model, we simulated *L* = 10^5^ causal sites which contribute to phenotypic variation, set *a* = 1, and choose *T* so as to attain the correct value of *p*_*T*_ . We set the mutation rate for causal sites to *µ* = 10^−7^. All causal sites are equally spaced along the chromosome. Between each pair of causal sites, there are 25 BGS sites, as well as a single neutral site (these neutral sites are used to empirically check the strength of the BGS effect, i.e. the *B* value, against the expectation). For most of our simulations, the recombination rate among adjacent sites is set to *r* = 2 × 10^−7^, and the mutation rate at BGS sites is set to *ν* = 2 × 10^−8^, to yield an expected *B* value of *e*^−0.2^ ≈ 0.82, approximately equal to the estimated human average. To ensure that all causal sites experience a similar background selection effect, we added 6.5 × 10^5^ negatively selected sites on each end of the chromosomes, such that the chromosome has a total map length of 0.8 Morgan. In the simulations where we vary the strength of the BGS effect, we do so by varying the mutation rate for BGS sites, *ν*, with the recombination rate held constant.

To compare traits with and without background selection, we first simulate a trait with the environmental variance tuned so that the heritability is *h*^2^ = 0.5 in the absence of background selection (i.e. setting *s* = 0 for all BGS sites). We then rerun the same simulation with background selection turned on (i.e. with *s* = −0.005 for all BGS sites), holding the environmental variance constant at the same value used in the no background selection case. In all cases, we set the fitness cost of being across the threshold at *C* = 0.2. An individual’s fitness is then computed as a product of their fitness effect due to the trait together with independent multiplicative contributions from the BGS sites.

To compare the single-effect-size trait model to the single-site results, we assign a scaled selection coefficient to each set of trait simulations. This quantity measures the average selection coefficient that causal variants experience in the trait model. For each choice of *p*^+^(*a*) in the simulations, we estimated the unscaled selection coefficient by regressing individual fitness against genotypes in the absence of background selection. This regression was performed for each causal site segregating in a given generation and repeated across multiple generations, with the average slope serving as our estimate. We then obtained the scaled selection coefficient by multiplying this estimate by 2*N* . The corresponding simulation run, conducted with the same *p*^+^(*a*) but with background selection included, was assigned the same scaled selection coefficient. In principle, in the small effect regime we could obtain the scaled selection coefficient analytically using Equation (13), but this method breaks down for large effects, so we use the empirical regression-based estimate across the full range.

#### Stabilizing Selection

Our simulations under stabilizing selection are similar to those under the threshold model, with a few key differences. Notably, to avoid the Bulmer effect in our simulations of stabilizing selection (see Supplementary section E and Figure S11), we had to increase the harmonic mean recombination rate between pairs of causal sites (Bulmer 1971; Bulmer 1974; Negm and Veller 2024; Veller and Coop 2024). In these simulations, we set *L* = 5.4 × 10^4^, and split the causal sites across 6 independent chromosomes, so that each chromosome has 9000 causal sites. Between each pair of adjacent causal sites on a given chromosome, there are 98 BGS sites and 1 neutral site. The recombination rate is set to *r* = 10^−6^ per base pair, while the mutation rate is *µ* = *ν* = 10^−7^ for both causal and BGS sites. We set the width of the stabilizing selection surface *ω*^2^ = 2*N* and vary the effect size for each site *a*.

To compare the single-effect-size trait model to the single-site results, for each choice of *a*, we measure the average heterozygosity of all the causal sites in the simulations without background selection over multiple generations and solved for the *τ* with Equation (23). We then multiply this estimate by 2*N* and assign it to the simulations with the same *a* with and without background selection.

#### Use of generative AI tools

We used generative AI tools, including Gemini 2.5 Pro and Gemini 3.0 Pro, as well as ChatGPT-5, to help draft and edit text in this manuscript, and to produce and edit code for plotting figures. All text and code drafted or edited in this manner was thoroughly checked and edited further before inclusion in the final manuscript. We take full responsibility for the contents of this manuscript.

## Resource

The documented code, including simulations, numerical solutions of the model, and figure-geneartion scripts, is publicly available at https://github.com/xinyli/BGS_msdb.git.

## Acknowledgements

We would like to thank John Novembre for many helpful conversations, and for reading and commenting extensively on several drafts of the manuscript. We would also like to thank members of the Berg, Novembre and Steinrücken labs for feedback on the work at various stages. This work was completed in part with resources provided by the University of Chicago’s Research Computing Center. XL was supported by NIH grant R35 GM149521 to John Novembre and R01 HG010773 to John Novembre and Xin He. JJB was supported by R35 GM151257.

## A. The frequency spectrum under background selection in a recombining background

**Table 1:**
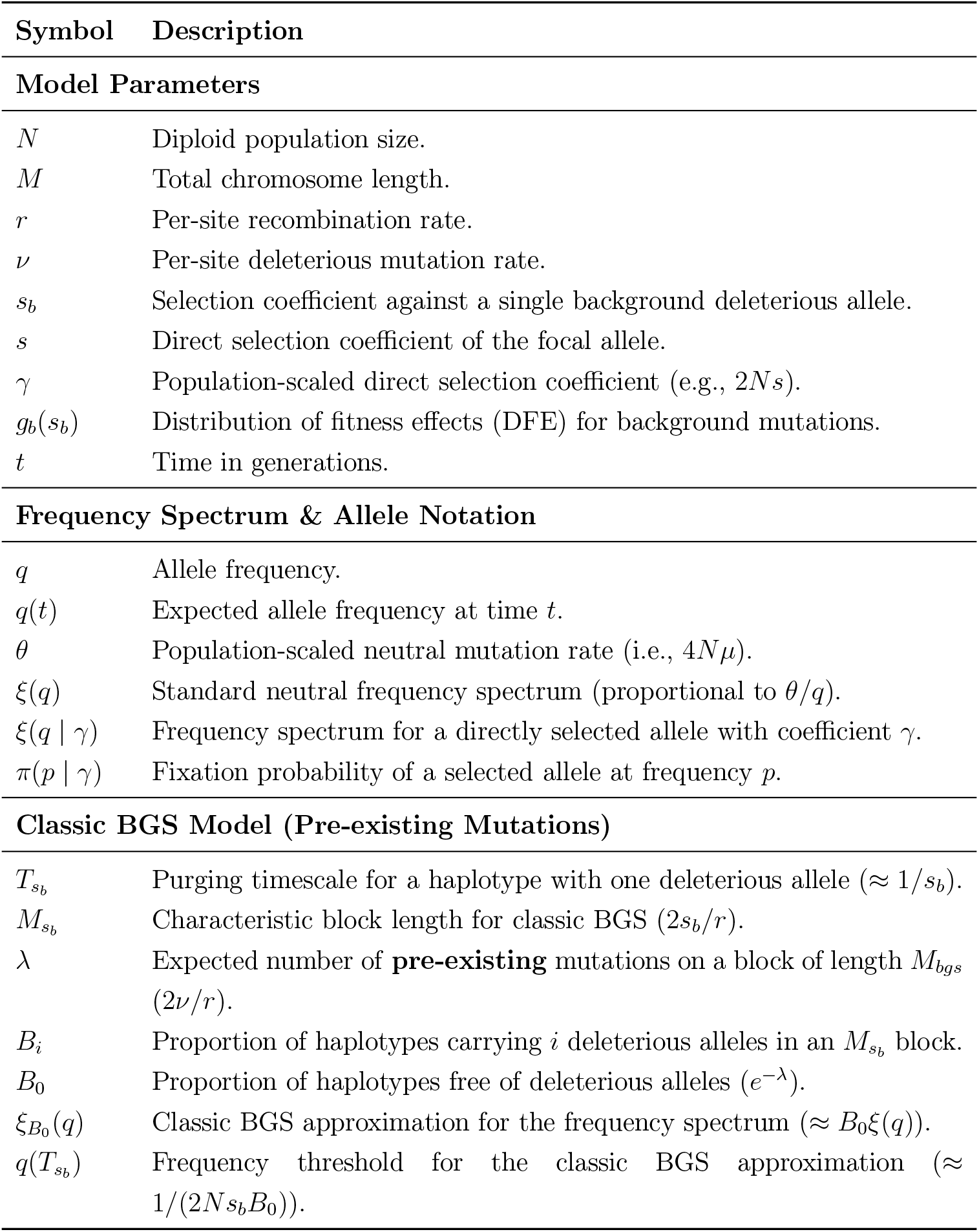
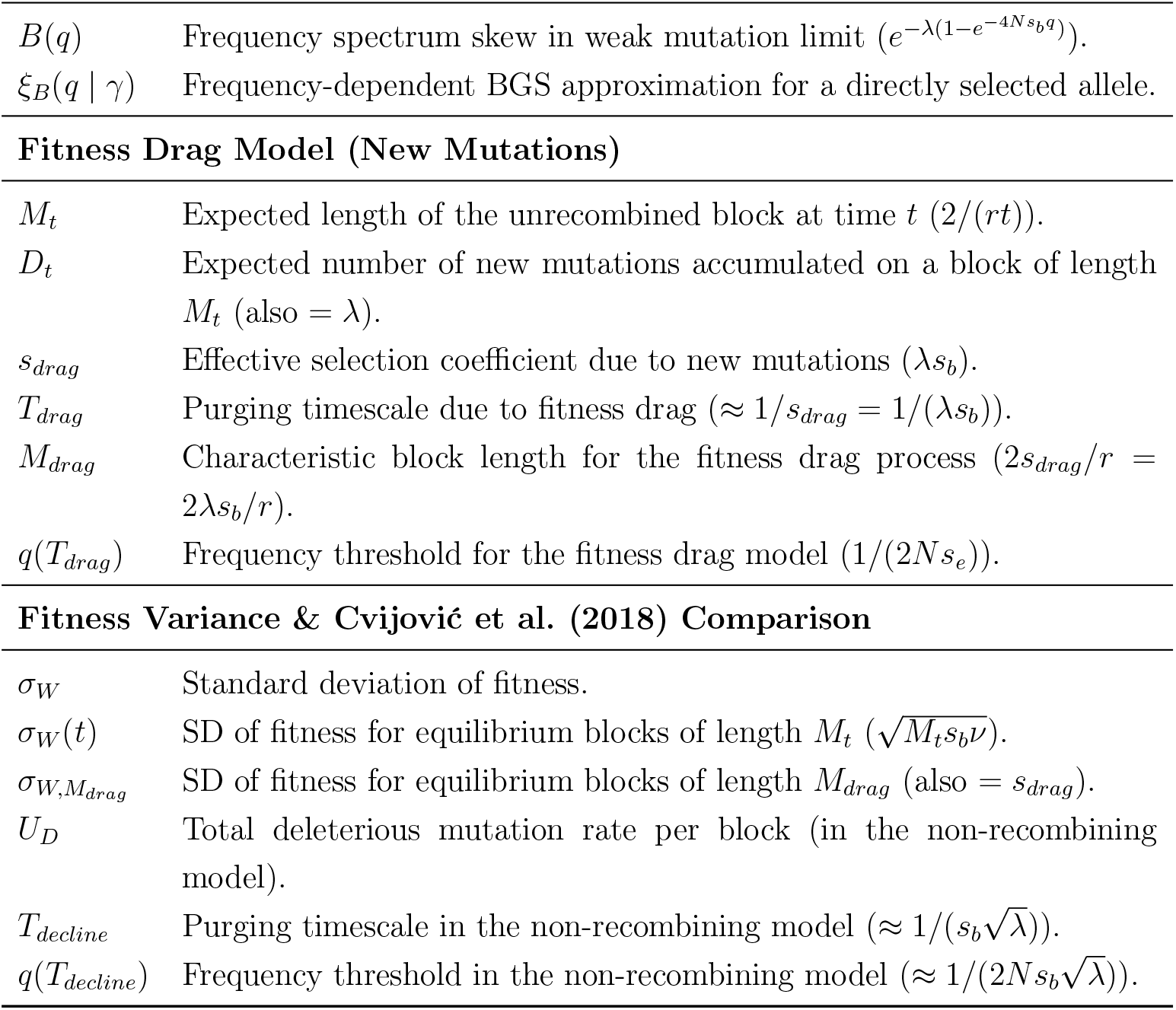
Notation table for Supplementary Text A.

### Classic background selection: removal due to pre-existing mutations

We consider a chromosome of length *M* recombining at rate *r* per site, on which deleterious alleles with selection coefficient *s*_*b*_ arise at rate *ν* per site. We are interested in the fate and frequency distribution of a focal allele that arises on a random chromosome in this population at time *t* = 0.

After *t* generations have passed, the distance (in each direction) from the focal allele to the first recombination event is exponentially distributed with rate *rt*. The length of the block that remains associated with the focal mutation without any intervening recombination is on expectation

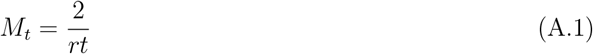

(the factor of 2 comes from the fact that there is an uninterrupted block with expected length 1*/rt* in each direction). A haplotype carrying a single deleterious allele is purged from the population in

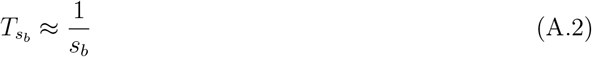

generations on expectation, so the expected length of a haplotype block over which selection discriminates between focal alleles linked to one vs. zero deleterious alleles is

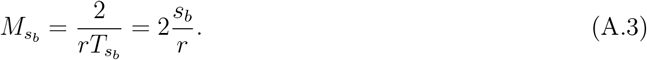

This defines a characteristic length scale in the classic model of background selection in a recombining background. At mutation-selection balance, the expected number of deleterious alleles in a window of this size is

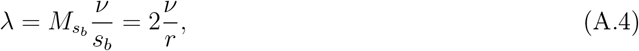

and the distribution is approximately Poisson, with the proportion of haplotypes carrying *i* deleterious alleles within this window given by

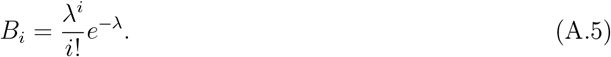

This result is the source of standard “reduced effective population size” approximation for a long recombining chromosome. In this approximation, only a neutral allele that arises on a length *M*_*bgs*_ haplotype free of deleterious mutations (which occurs with probability *B*_0_ = *e*^−*λ*^) has any hope of being seen at appreciable frequency in the population. Therefore, while the neutral frequency spectrum in the absence of background selection would be expected to take the form

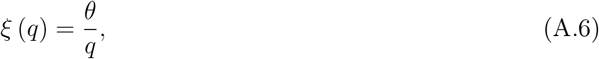

the classic background selection approximation implies a systematic reduction by a factor of *B*_0_ in the number of alleles reaching any given frequency

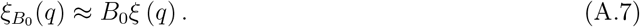

Because the effect of selection is not instantaneous, this approximation is only valid above a certain critical frequency. To determine what this frequency is, consider the transit of a focal neutral allele linked to a haplotype block carrying a single deleterious allele. This haplotype block originated from within the population of mutation free blocks, and all other blocks carrying at least one mutation will soon be extinct, so we can think of it as evolving within a population of reduced effective size *NB*_0_.

As we discussed above, the allele will be eliminated by selection on a timescale of 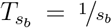 generations, and so will be prohibited from reaching frequencies that cannot be reached via genetic drift in that amount of time. For small values of *t*, the expected frequency of a neutral allele in a diploid population of size *NB*_0_ conditional on not yet having been lost is approximately *q* (*t*) ≈ *t/*2*N B*_0_. Therefore, haplotypes carrying at least one deleterious allele are limited to frequencies no greater than approximately

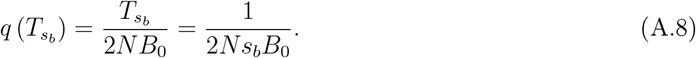

Equation (A.8) thus gives an approximate threshold for the frequency at which Equation (A.7) becomes valid. This is the the threshold suggested by Cvijović et al. 2018, provided that one replaces 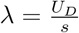 in their haploid model of a non-recombining block with 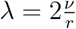 in our diploid model of a recombining chromosome.

### Length- and time-scale of removal due to selective drag of new mutations

The classic background selection model considered above assumes that the most relevant process is the removal of the haplotype that the focal allele arises on, due to selection against mutations that are already present there when the allele arises. If the deleterious mutation rate is high relative to the recombination rate, then it could also be removed due to selection against deleterious mutations accumulating after it arises. After *t* generations have passed, the haplotype associated with the deleterious allele has expected length *M*_*t*_ ≈ 2*/rt* (see Eq. (A.1)), and so on expectation has accumulated

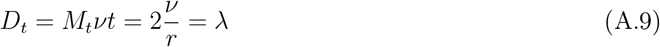

additional deleterious alleles since it arose. It is notable that this expectation is independent of time, as the deleterious alleles that are separated by recombination are replaced by new deleterious mutations arising on the (on expectation, slightly shortened) block that remains associated. While their expectations are numerically equal (*D*_*t*_ = *λ*), it is important to distinguish these two quantities conceptually: *D*_*t*_ is the expected number of new mutations accumulated on a shrinking, age-*t* block, whereas *λ* is the expected number of pre-existing mutations found on an equilibrium block of length *M*_*bgs*_.

This time-independence represents a key distinction from the non-recombining model of Cvijović et al. 2018, in which the absence of recombination means that the number of new mutations accumulates linearly with time (i.e. *D*_*t*_ ∝ *t*). In that case, the process is one of steady fitness decline as the haplotype becomes progressively loaded with mutations. In the recombining model, the balance between recombination and mutation establishes a constant, time-independent expected mutational load, *D*_*t*_ = *λ*, for the block associated with the focal allele. (We say more about the similarities and distinctions between the two models below.)

Each recombination event generates a new set of associations with deleterious alleles at distances greater than the expected 1*/rt* distance to the edge of the block. However, these new associations are samples from the equilibrium distribution, and therefore have contributions that are on average equal to the population mean, and thus have no net effect on the mean fitness of the focal block. The result is that in the early generations before selection has had a chance to shape the trajectory of our focal allele, its expected fitness drag due to new mutations is constant:

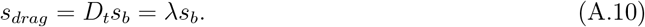

This suggests that a typical allele at the focal sites should be removed due to this immediately accumulated selective drag on a timescale of

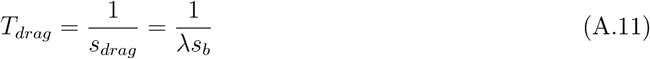

generations, and that the expected length of the haplotype block that is removed with it is

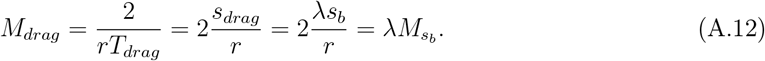

This process of selection against newly mutations arising on the background of the focal allele will be important for frequencies less than

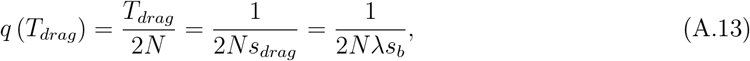

but becomes unimportant above this threshold because any alleles exceeding this frequency are likely outliers that have accumulated fewer deleterious mutations than average.

As we indicated above, this process is only important to the dynamics if recombination is weak relative to mutation, in which case the timescale of removal due to the accumulation of deleterious mutations is faster than the timescale on which mutation-free haplotypes are preserved, i.e. 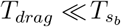 . Intuitively, this condition translates into a requirement that *λ* ≫ 1.

More precisely, we might think that in order for the effects of the immediate fitness drag to be important, the threshold frequency at which this fitness drag has substantially reshaped the frequency spectrum must be below the threshold frequency at which the classic background approximation begins to apply, i.e. 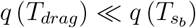. This is equivalent to the condition that 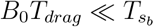 . This leads to the more restrictive condition *λ* ≫ *e*^−*λ*^ on the value of *λ* at which the fitness drag becomes important. This is solved by *λ* ≫ *W*_0_ (1) ≈ 1*/*2, i.e. 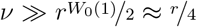, where *W*_0_ is the main branch of the Lambert W function and, very roughly, *W*_0_(1) ≈ 1*/*2. This condition more closely matches our observations from simulations that the fitness drag begins to impact the frequency spectrum roughly when *λ >* 1*/*2 (Figure S1).

#### Comparison to Cvijović et al. (2018) non-recombining model

This “fitness drag” process provides a close parallel to the “fitness decline” process in the non-recombining model of Cvijović et al. 2018. In that model, *λ* is defined as 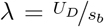, where *U*_*D*_ is the total deleterious mutation rate for the entire non-recombining block. For an equivalent value of *λ*, both models predict the same overall reduction in effective population size (*B*_0_ = *e*^−*λ*^). Furthermore, if *s*_*b*_ is also held constant, the upper frequency threshold 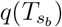—which defines the frequency below which the classic BGS approximation of Equation (A.7) breaks down—is also the same. This equivalence follows directly from the validity of the “effectively non-recombining block” approximation, which simply converts the recombining problem into the corresponding non-recombining one for alleles above the relevant threshold.

The most significant difference between the two models lies in their predictions regarding the impact of selection on alleles at very low frequencies. Notably, because the “block length” is fixed in the non-recombining model, there is no contribution from the accumulation of deleterious alleles at distances beyond the characteristic block length. Instead, the block must first decline in fitness by accumulating deleterious mutations. As a result, we speak of a process of “fitness decline” and its consequences, rather than the “fitness drag” process described above for the recombining model.

In the non-recombining model, these accumulated mutations become important once the fitness of the block has declined by an amount equal to the equilibrium standard deviation in fitness, 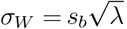. At this point, the fitness deficit relative to the rest of the population is substantial enough that selection can effectively remove the allele.

Ignoring the impact of selection on short timescales, fitness declines linearly with time in the non-recombining model (i.e., *s*_*e*_(*t*) ≈ (*U*_*D*_*t*)*s*_*b*_). Setting the accumulated load (*U*_*D*_*t*)*s*_*b*_ equal to the equilibrium standard deviation 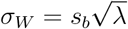 and solving for *t*, the timescale on which an allele declines sufficiently in fitness to be removed by selection is

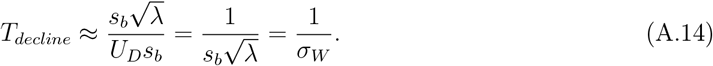

Consequently, in the non-recombining model, we expect that for frequencies below a threshold of

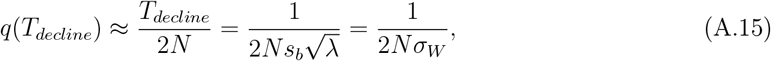

selection has not yet had an opportunity to act, so the frequency spectrum should be approximately neutral. Once an allele reaches this frequency, however, it is likely to have accumulated a fitness deficit of approximately *s*_*e*_ (*T*_*decline*_) = *σ*_*W*_, and will therefore be eliminated on a timescale of roughly *T*_*decline*_ = 1*/σ*_*W*_ additional generations. During this time, the allele’s frequency can increase by at most another factor of roughly 2. Thus, equation (A.15) provides the correct order-of-magnitude limit on the frequency at which the accumulation of new mutations dominates the dynamics.

Notably, the equivalent threshold in the recombining model, *q* (*T*_*drag*_) = 1*/*2*N s*_*b*_*λ* (Eq., (A.13)), is lower by a factor of 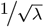. This difference reflects the difference in dynamics between the two models which can also be understood in terms of an appropriate standard deviation of fitness. Specifically, in the recombining model, the drag *s*_*drag*_ is equal to the equilibrium fitness standard deviation of blocks with length *M*_*drag*_ (i.e., blocks of the characteristic length eliminated by this process). The equilibrium distribution of pre-existing mutations on a block of length *M*_*drag*_ is Poisson with mean *M*_*drag*_(*ν/s*_*b*_) = (2*λs*_*b*_*/r*)(*ν/s*_*b*_) = *λ*^2^. The standard deviation of fitness among blocks of this length is therefore:

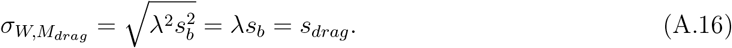

This equality provides a dynamic interpretation of the removal timescale, *T*_*drag*_. When an allele is young (*t* ≪ *T*_*drag*_), its associated block is long (*M*_*t*_ ≫ *M*_*drag*_). The background standard deviation of fitness for very young blocks is consequently very large 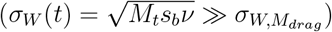, and the allele’s constant fitness drag is small relative to this background variation. As the allele ages and *t* approaches *T*_*drag*_, its associated block shrinks toward length *M*_*drag*_, and the background fitness variance shrinks with it. When the block reaches length *M*_*drag*_, the background variance shrinks to equal the fitness drag 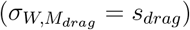, and it is around this time that the allele is likely to be purged.

This difference in mechanism—the accumulation of fitness deficit versus a constant fitness drag—arises because, in the recombining model, deleterious mutations far beyond the characteristic block length of classic BGS exert an immediate influence. This extended range accelerates the onset of strong selection: the average fitness deficit becomes visible to selection on a timescale shorter by a factor of

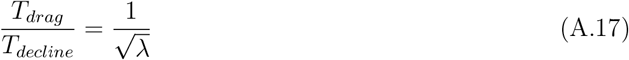

compared to the equivalent non-recombining model, and consequently that BGS should skew the frequency spectrum at frequencies that are also lower by a factor of

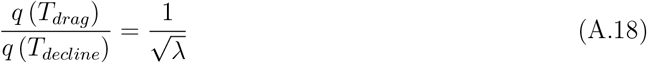

This perspective on the recombining model is somewhat at odds with the idea that the allele is under strong, constant selection immediately. For early times *t* ≪ *T*_*drag*_, when the drag is much smaller than the background variance (*s*_*drag*_ ≪ *σ*_*W*_ (*t*)), the argument made by Cvijović et al. (2018) in the context of the non-recombining model would suggest that the allele should behave neutrally at first due to the noise of the background. This tension is in some part real, and reflects the fact that all of these are coarse approximations for an extremely complex process. However, this tension is partially resolved by the fact that the argument from Cvijović et al. (2018) is a bit too strong, and also partially by the difference in dynamics between the two models.

In the non-recombining model, the fitness deficit grows linearly relative to a fixed standard deviation, meaning the ratio also scales linearly with time:

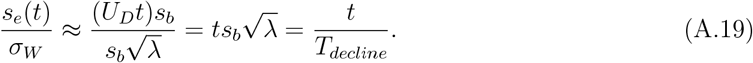

Consequently, the time-averaged value of *s*_*e*_(*t*)*/σ*_*W*_ over the interval [0, *T*_*decline*_]—the period during which the allele’s fitness deficit increases from 0 to *σ*_*W*_ —is 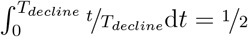 . This suggests that if one wishes to approximate the early dynamics using a constant selection coefficient, 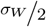 may be a more useful choice than 0.

In contrast, in the recombining model, the background fitness standard deviation shrinks with 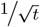, while the fitness deficit remains constant. Consequently, the ratio scales with the square root of time:

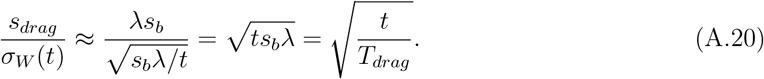

The time-average of 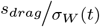 over the interval [0, *T*_*drag*_] is therefore 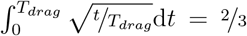 . This reflects the fact that the ratio of the fitness deficit to the background standard deviation is higher on average during the early phases of the allele’s trajectory than in the non-recombining case (Figure S4). This suggests that to approximate the low-frequency dynamics of the recombining model using a single constant selection coefficient, one should use 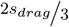, thereby discounting for the impact of background fitness variance in the earliest generations.

These arguments suggest that the frequency spectrum is more strongly distorted in recombining models than in equivalent non-recombining ones, not only because the focal allele becomes “visible” to selection faster, but also because it spends more time near this visibility threshold before reaching it. While these arguments are heuristic, the central conclusion is deeply intuitive: the suppression of the frequency spectrum extends to lower frequencies in large recombining genomes than predicted by non-recombining models with equivalent reductions in effective size, due to the action in early generations of deleterious mutations at recombination distances greater than the characteristic length-scale of the classical recombining BGS model.

### The frequency spectrum at low frequencies

#### The strong recombination/weak mutation regime

##### Neutral alleles

The arguments in the preceding sections allow us to understand the shape of the frequency spectrum for neutral alleles at very low and very high frequencies. Roughly, when *λ* ≪ 1*/*2 (or equivalently, when *B*_0_ ≫ 0.6) recombination is too fast relative to mutation for the process of fitness decline to be relevant (i.e. 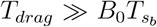), and so the frequency spectrum is shaped entirely by the effect of selection acting against deleterious alleles that are already present when the focal neutral allele arises. Based on the approximations given above, this suggests that

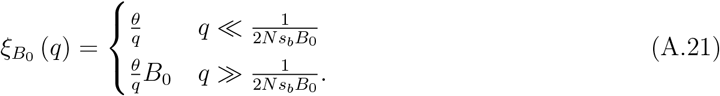

We can push this perspective further and obtain an approximation that is valid across all frequencies by taking advantage of the assumption that the focal allele is eliminated by deleterious alleles already present on its background when it arises. To this end, consider that with probability *B*_*i*_ the focal allele arises on a haplotype that carries *i* deleterious alleles within a distance of 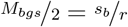 in either direction. Provided that carrying *any* deleterious alleles on a haplotype of this length is rare in general (consistent with the assumption that *B*_0_ ≫ 0.6), then standard diffusion results tell us a neutral allele on a haplotype with *i* deleterious alleles will spend approximately 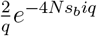 generations at frequency *q*. We can therefore approximate the frequency spectrum across the full range of frequencies as

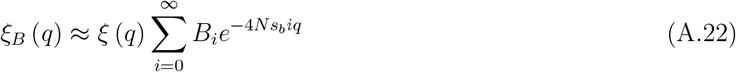

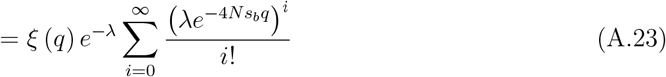

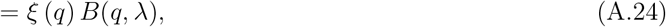

where

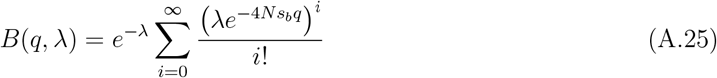

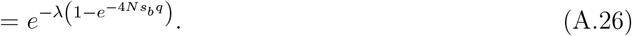

is a frequency dependent B value, the form of which follows from the fact that the infinite sum in Equation (A.25) is the full Taylor series representation of the function 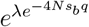 expanded around the point 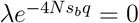. We note that the sum in equation H3 of Cvijović et al. 2018 is essentially identical to our Equation (A.22) above, although they evaluate it only to zeroth order, i.e., *ξ*_*B*_ (*q*) = *ξ* (*q*)+ 𝒪 (*λ*).

##### A distribution of background fitness effects

This frequency dependent B value can easily be generalized for a distribution of fitness effects among the sites generating background selection. Suppose that sites responsible for background selection draw their selection coefficients from some distribution, *g*_*b*_ (*s*_*b*_), and suppose that the largest selection coefficients sampled from this distribution still conform to the simplifying assumptions that *s*_*b*_ ≪ *Mr* and 2*Ns*_*b*_ ≫ 1.

Under these assumptions, the number of linked deleterious mutations is Poisson distributed with mean *λ*, and the selection coefficient of each mutation is drawn independently from *g*_*b*_(*s*_*b*_). Since the effect of each mutation on the sojourn time is multiplicative (multiplying the neutral time by 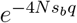), the total frequency dependent B value is the expectation of the product of these effects, yielding:

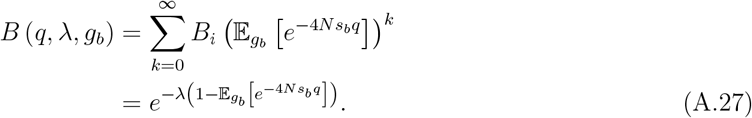

where 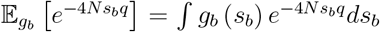. Note that this derivation appears to neglect the fact that the characteristic length scale depends on the selection coefficient. However, while we might expect this dependency to complicate the strong mutation limit, it is irrelevant in the weak mutation limit. In this regime, a haplotype is typically eliminated by a single deleterious allele; consequently, the effective length scale is determined solely by the mutation responsible for the elimination. The result therefore correctly averages over the relevant distribution of length scales.

##### Directly selected alleles

When the focal allele undergoing background selection is itself directly selected, we can find useful approximations by straightforwardly extending the results above under neutrality. First, consider the expected frequency spectrum of a deleterious allele in the absence of linked selection, which is

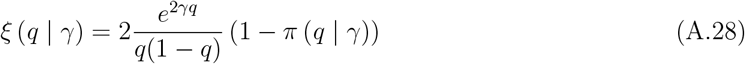

where

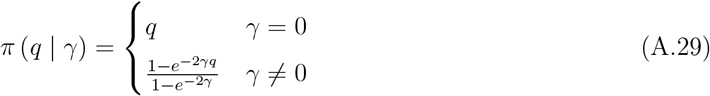

is the fixation probability of an allele at frequency *q* with scaled selection coefficient *γ*.

In Charlesworth (1994)’s classical approximation for weakly selected alleles (i.e. *γ* ∼ 1), we simply ignore all variants arising on backgrounds other than the mutation free background, i.e.

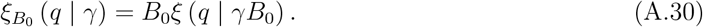

This approximation extends the “scaled mutation rate” effect of background selection by including a “scaled selection coefficient” effect. The argument for this approximation relies on a separation of timescales between background selection (which acts quickly), and direct selection (which is assumed to act slowly). Alleles reaching frequencies greater than 1*/*2*N s*_*b*_*B*_0_ must have originated on backgrounds with *i* = 0 deleterious alleles. Once these alleles reach appreciable frequency, new deleterious mutations occurring on their background spread the focal allele across all fitness classes, but because backgrounds with *i >* 0 deleterious alleles are always quickly removed, only the population of size *NB*_0_ is relevant to the evolution of the focal allele. As a result, the balance between drift and selection over the longer timescale on which (weak) direct selection acts is captured by a rescaled version of the population scaled selection coefficient: *γB*_0_.

Combining this classic separation of timescales argument with the frequency dependent B value argument we articulated above, we can approximate the full frequency spectrum of directly selected alleles under weak selection as

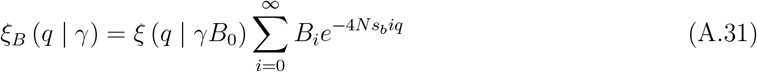

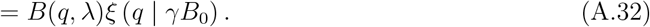

Because *B*_0_ = *e*^−*λ*^ regardless of the value of *s*_*b*_ (provided that *s*_*b*_ ≪ *Mr*), this approximation can be straightforwardly extended to the case with a distribution of background fitness effects using Equation (A.27).

Alternatively, when the strength of direct selection is similar to that of background selection, this separation of timescales does not apply. Direct and background selection act on the same timescale, leading to:

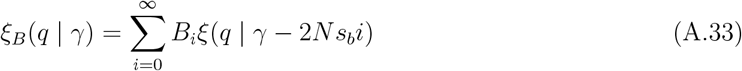

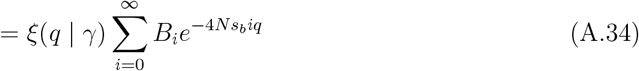

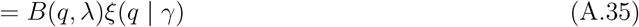

#### The strong mutation/weak recombination regime

When recombination is weak relative to mutation (i.e., *λ* ≫ 1*/*2), the frequency spectrum is shaped differently. As established, this condition implies 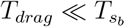, creating two distinct selective thresholds, 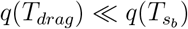, which define three frequency regimes. In the lowest frequency regime, *q* ≪ *q*(*T*_*drag*_), the frequency spectrum is shaped by the emergence of the time-independent, constant expected fitness drag against the shrinking background fitness variation. In the highest frequency regime, 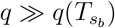, all alleles originated on the fittest background, and the frequency spectrum is well approximated by the classic rescaled effective population size. For the intermediate regime, where 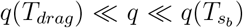, Cvijović et al. (2018) provide an approximation in the non-recombining model for the form of the frequency spectrum. Here, for completeness, we use heuristic arguments to recapitulate their results in the context of the recombining model.

The key to this regime is that our constant *s*_*drag*_ approximation, while useful for describing the behavior at frequencies below the lower threshold *q*(*T*_*drag*_), breaks down at frequencies above it. Specifically, *s*_*drag*_ = *λs*_*b*_ represents the expected fitness drag for an allele that has not yet persisted long enough for selection to significantly impact its trajectory. For an allele to reach a frequency *q* ≫ *q*(*T*_*drag*_), it must have been lucky, arising on a background with *i* ≪ *λ* deleterious alleles.

Consequently, the dynamics of the intermediate regime are governed by the “classic” BGS process.

While this is the same process as in the *λ* ≪ 1 case, the required mathematical approximation is different. When *λ* ≪ 1, the variance of the distribution of *i* is low, and the fittest class (i.e., *i* = 0) is essentially the same as the population mean. In contrast, when *λ* ≫ 1, the variance is high, and the lucky alleles that survive into the intermediate regime are extreme outliers, far fitter than the population mean (i.e., *i* ≪ *λ*).

When *λ* ≫ 1, the probability of arising on a background with *i* deleterious alleles declines quickly as *i* → 0. It follows that the set of backgrounds reaching frequency *q* will be dominated by the least fit backgrounds (i.e., those with the highest value of *i*) that are nonetheless capable of reaching frequency *q*. We can heuristically derive this least fit class, *i*(*q*), via an extreme value argument. To reach a frequency *q*, an allele must arise on one of the fittest “available” backgrounds. We can estimate the number of “available” backgrounds as 2*Ns*_*b*_*q* (the number of individuals at that frequency, 2*Nq*, divided by the selection timescale, 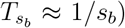. The fittest available background class, *i*, is the one that solves the extreme value condition:

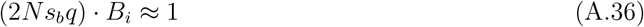

where 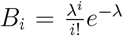 is the probability of arising on a background with *i* deleterious alleles within the characteristic block of length *M*_*bgs*_ (Eq. (A.5)). Taking the log of both sides and applying Stirling’s approximation (ln(*i*!) ≈ *i* ln *i* − *i*) gives:

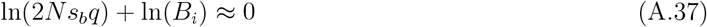

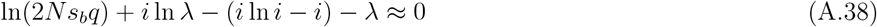

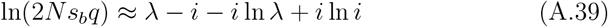

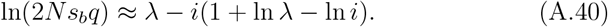

Assuming that the ln *λ* term dominates, we can approximate 1 + ln *λ* − ln *i* ≈ ln *λ* (this holds when *λ* is large and *i* is effectively intermediate—i.e., 1 ≪ *i* ≪ *λ*). This allows us to solve for *i* as a function of *q*:

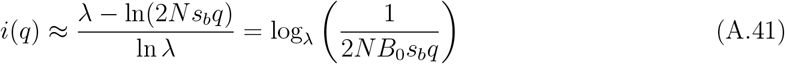

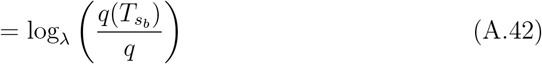

This *i*(*q*) represents a “self-consistency” condition (equivalent to *k*_*c*_(*f*) + 1 in Cvijović et al. 2018), which shows that the dominant fitness class among backgrounds reaching frequency *q* declines loga-rithmically with increasing frequency. The alternative form for *i* (*q*) in Equation (A.42) follows from the definition 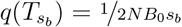. This form emphasizes that as *q* increases, the allele must have been increasingly lucky to survive: to reach higher frequencies, it must have arisen on a background with fewer deleterious alleles, until eventually, to reach frequency 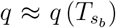, the allele must have arisen on a background with *i* = 0 (by which point this approximation has broken down).

The expected number of generations an allele spends at frequency 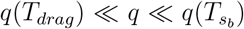 is thus approximately

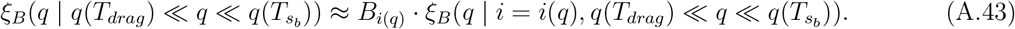

We already assumed above that *B*_*i*(*q*)_ ≈ 1*/*2*N s*_*b*_*q* (Eq. (A.36)). Given that the allele arises in this class, the amount of time spent at frequency *q* has the approximate form

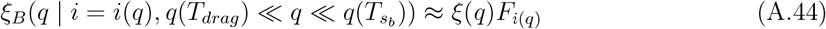

where *ξ*(*q*) = *θ/q* is the standard neutral spectrum and *F*_*i*(*q*)_ is a factor that accounts for the suppression of this neutral frequency spectrum by selection in the time since the allele arose. Given the fundamental timescale, 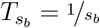, of the background selection process, *F*_*i*(*q*)_ can be approximated as

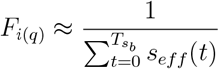

where *s*_*eff*_ (*t*) is the effective selection coefficient experienced by the allele in generation *t*. Notably, *s*_*eff*_ (0) = *s*_*b*_*i*(*q*), but the allele will experience a range of selective environments over its transit through the population, due to a combination of mutation and recombination events shuffling it across the distribution of fitness backgrounds. This process is complicated, so we might naturally want to make an approximation 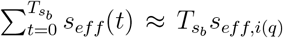, for some appropriate average effective selectioncoefficient, *s*_*eff,i*(*q*)_. Most backgrounds are less fit than the *i*(*q*) background on which it arose, which suggests that *s*_*eff,i*(*q*)_ *> s*_*eff*_ (0). Alternatively, in order to be found at frequency *q*, the allele cannot have spent too much time on much less fit backgrounds, or it would have been purged. When one conducts a rigorous analysis of the dynamics (as Cvijović et al. 2018, do for the non-recombining case), one finds that 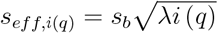, which is the geometric mean of the allele’s initial selection coefficient *s*_*eff*_ (0) = *s*_*b*_*i*(*q*), and the one that it would experience on average if it were sampling the distribution completely at random, i.e., *s*_*b*_*λ*, and therefore nicely satisfies both of these criteria.

It follows that

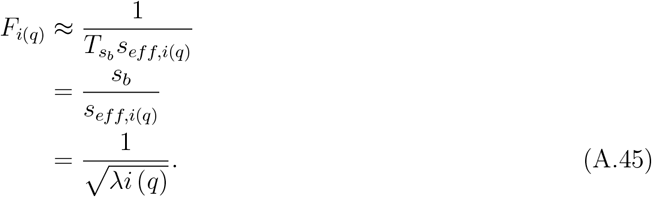

Thus, putting together Eqs. (A.41)-(A.45), we have

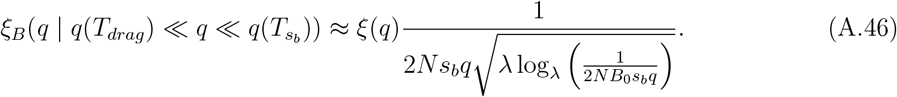

Combining this with the heuristic 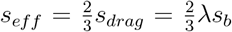 approximation we derived above for the *q* ≪ *q*(*T*_*drag*_) regime, this allows us to write an approximation for the full spectrum for neutral alleles in this regime as a three-part piecewise function, analogous to Equation I21 in Cvijović et al. 2018:

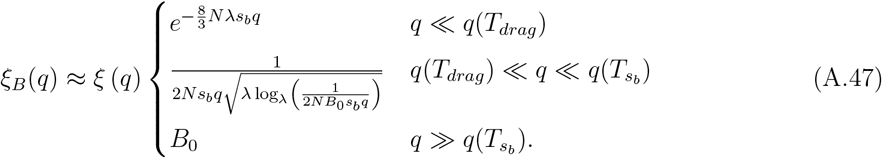

Notably, this approximation neglects the deterministic sweep-like behavior described by Cvijović et al. (2018) at very high frequencies. We expect that an effectively non-recombining block approximation could be derived for this regime as well.

## B. Derivations Supporting the Exponential Fitness Model

### Selection coefficient

To begin, we divide an individual’s phenotype into a contribution from the focal site *ℓ*, which is *a*_*ℓ*_*g*_*ℓ*_ (where *g*_*ℓ*_ ∈ {0, 1, 2} is the genotype at site *ℓ*), and the contribution from all other sites, which we write as *Z*_−*ℓ*_. Writing *f* (*Z*_−*ℓ*_) for the density on this background contribution, the expected fitness of an individual with genotype *g*_*ℓ*_ is exactly

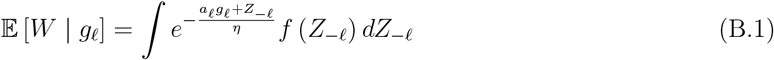

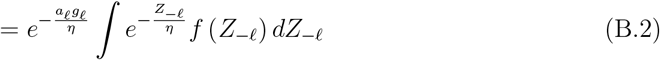

The ability to factor the fitness effect of the focal site *ℓ* out of the total effect is a special property of the exponential model, which allows us to rescale the relative fitnesses so that independent of any details about the background, *Z*_−*ℓ*_, we can write

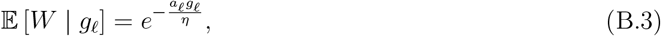

without any loss of generality or approximation.

The expected change in frequency for an allele with effect *a* and frequency *x* can be written as

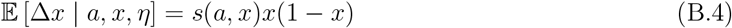

where

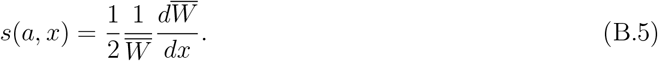

is the marginal effect of the allele on fitness. The mean fitness is equal to

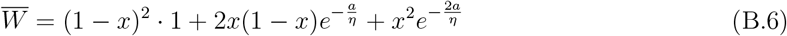

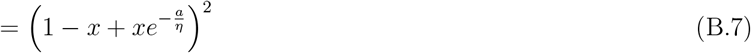

so that

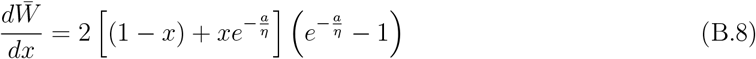

and therefore

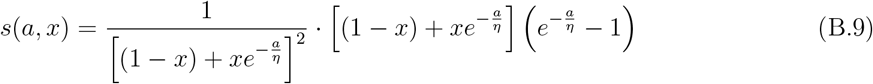

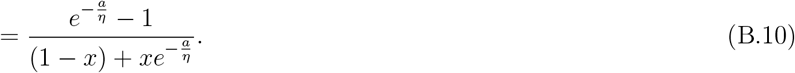

Equation (B.10) is still exact. Assuming that 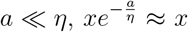, and 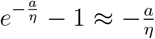, so we can make the approximation

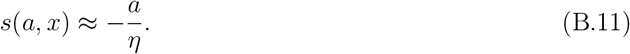

### Impact of BGS on the Mean Phenotype and Fitness Load

Here we derive the effect of background selection on the mean phenotype and the corresponding fitness load. The mean phenotype is given by the sum of contributions from sites fixed for the trait-increasing allele. At mutation-selection-drift balance, the fraction of sites fixed for the trait-increasing (deleterious) allele with scaled selection coefficient *γ* is

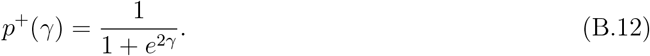

In the presence of background selection, the effective population size is reduced by a factor *B*, rescaling the selection coefficient to *γB*. The change in the fraction of fixed deleterious alleles is therefore

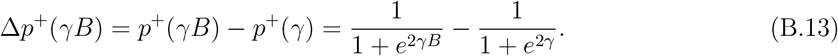

The shift in the mean phenotype is the sum of the effects of these additional fixed alleles across all *L* loci:

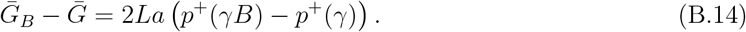

To compare this shift to the scale of the standing genetic variation, we normalize by the genetic standard deviation 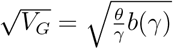, where *b*(*γ*) = tanh(*γ*):

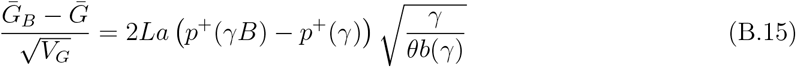

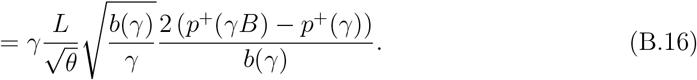

For weakly selected sites (*γ* ≪ 1), we can approximate 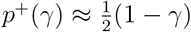 and *b*(*γ*) ≈ *γ*. Substituting these approximations yields:

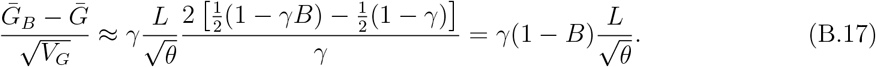

Since 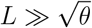, this standardized shift is substantial (≫ 1) even for moderate background selection.

This shift in the mean phenotype results in a decline in mean fitness. The ratio of mean fitness with BGS to mean fitness without BGS is:

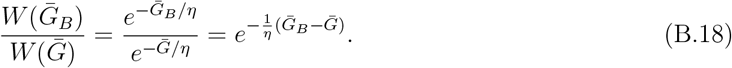

Substituting the expression for the shift in the mean:

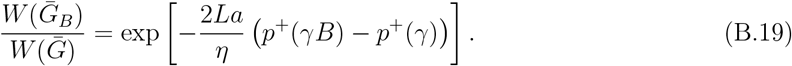

Using the relation 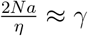 (so 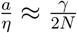), this becomes:

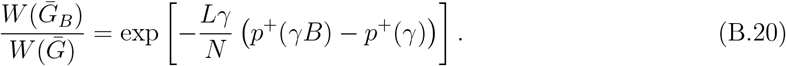

For weak selection (*γ* ≪ 1), we use the approximation 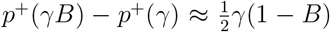. Furthermore, note that *p*^+^(*γ*) ≈ 1*/*2 and *p*^−^(*γ*) ≈ 1*/*2, so 4*p*^+^(*γ*)*p*^−^(*γ*) ≈ 1. Thus, we can write:

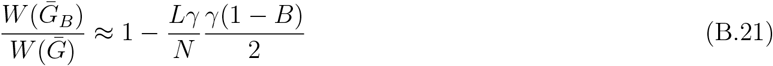

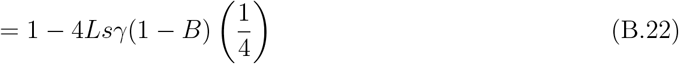

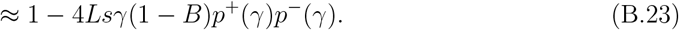

This result shows that for weakly selected sites, background selection drives a reduction in mean fitness proportional to the strength of selection, the total mutation rate, and the reduction in effective population size.

## C. Changes in disease prevalence in the liability threshold model due to BGS

### Derivation

Let *V*_*E*_ environmental variance, and let *V*_*G*_ be the genetic variance in the absence of background selection. The heritability in the absence of background selection is then

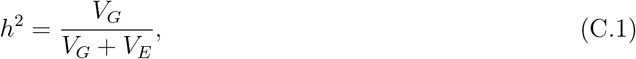

and the standardized threshold density is

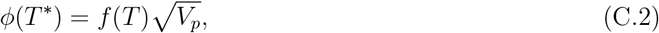

where 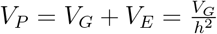 is the total variance.

In the presence of background selection, the genetic variance is reduced to *V*_*G,B*_ = *BV*_*G*_, and the phenotypic variance to *V*_*P,B*_ = *V*_*G,B*_ + *V*_*E*_, so the heritability becomes

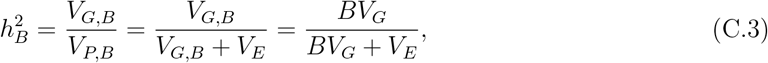

and we can write the standardized threshold density in the presence of background selection in terms of original quantities in its absence as

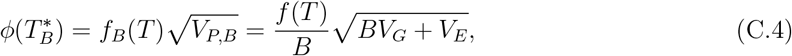

where 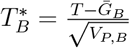 .

Now, we can eliminate the *V*_*E*_ by solving Equation (C.1) for 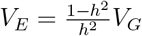 and substituting to get

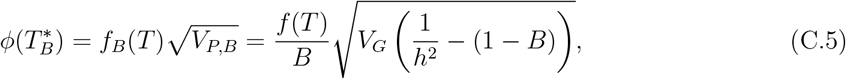

which is now written entirely in terms of counterfactual quantities from the equilibrium in the absence of background selection, and the B value. The increase in the standardized threshold density due to background selection is thus

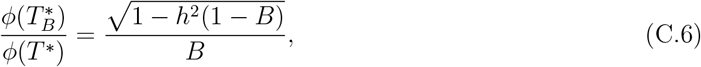

Translating this into a statement about the effect on the prevalence requires that we make some assumption about the relationship between the standardized threshold density and the prevalence. The standard assumption is that the distribution of liability is Normal, in which case the two expressions for the standardized threshold density above can be plugged into standard normal expressions to obtain the prevalence in each case. However, Berg et al. (2025) argued that when the prevalence is low, the relationship between the standardized threshold density and the prevalence is very roughly linear. It follows that

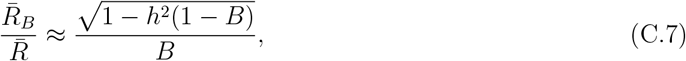

where 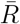 is the prevalence without BGS and 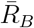 the prevalence with BGS (i.e. equation (17) in the main text). Berg et al. (2025) also showed that if large effect sites make a substantial contribution to genetic variance in liability, the liability distribution may become skewed. However, due to the polygenic background, the far tail of the distribution will still be roughly Gaussian in shape, and it is the rapid Gaussian decay of the tail that is responsible for the fat that the change in threshold density roughly predicts the change in prevalence (because most of the density in the tail that is beyond the threshold is nonetheless bery close to it), and so we expect equation (C.7) to hold, at least approximately, even if there is some skew in the distribution.

### Negligible impact of frequency spectrum skew

We also used simulations to explore whether this skew in the frequency spectrum impacts the disease prevalence. To do this, we varied the selection coefficients at the background selection sites while holding the mutation and recombination rates (and therefore the local effective size reduction) constant. We found that while the reduction in heterozygosity remained constant with increasing selection coefficients, the mean derived allele frequency among rare variants (*AF <* 0.1) rose, as expected. However, this increase in rare allele frequency had no discernible impact on the prevalence (Figure S12).

## D. Solving the Two Effect-size Model with Background Selection

Here, we detail the method for solving the two-effect-size liability threshold model in the presence of background selection. Berg et al. (2025) describes the method for solving this model in the absence of background selection in their Supplementary Section S7. Here, for completeness, we largely reproduced that description, with the addition of the BGS effect (as well as a few minor differences in notation).

In our two-effect model, we assume that a fraction *p*_*S*_ = 1 −*p*_*L*_ of sites have small effects, while the remaining fraction *p*_*L*_ have large effects. We model the distribution of liability in the population as a convolution of two distributions. The first is a Normal component with variance *V*_*G,B,S*_ + *V*_*E*_, where *V*_*G,B,S*_ is the variance from small effect sites (in the presence of background selection), and *V*_*E*_ is the environmental variance. The second is a Poisson distribution on the number of large effect alleles that an individual carries. This Poisson distribution has mean

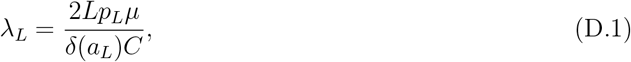

where

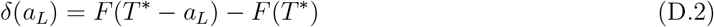

is the risk effect of the large effect sites, 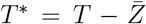 is the distance between the mean and the threshold,

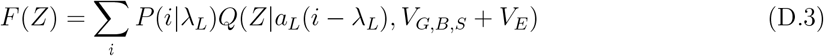

is the probability that an individual’s liability exceeds a value of *Z*, where *Q*(*Z*|*u, σ*^2^) = 1−Φ(*Z*|*u, σ*^2^) is the complementary CDF of a Normal distribution with mean *u* and variance *σ*^2^, and *P* (*i*|*λ*_*L*_) is the probability that a Poisson random variable with mean *λ*_*L*_ takes a value of *i*. The density on total liability, in turn, is

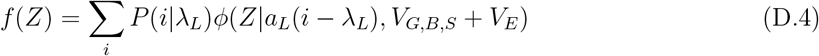

where *ϕ*(*Z*|*u, σ*^2^) is the Normal PDF.

We solve two-effect model in the following way. First, we can solve for the fraction of sites fixed for the liability increasing allele at small effect sites. Because all large effect sites are fixed for the liability decreasing allele, this fraction is equal to

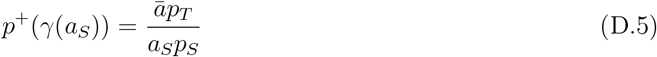

where

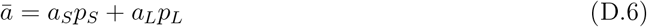

is the mean effect size. The scaled selection coefficient of the small effect sites is determined entirely by the threshold induced fixation asymmetry and the resulting long-term fixation dynamics, and is therefore given by

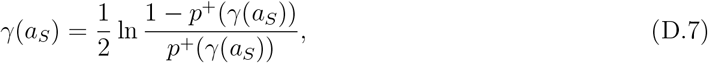

and the threshold density by

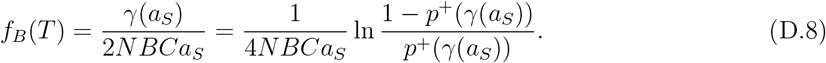

The genetic variance due to small effect sites is

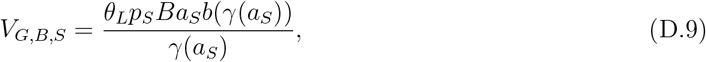

where *b*(*γ*(*a*_*S*_)) = 1 − 2*p*^+^(*γ*(*a*_*S*_)) is the degree of mutational asymmetry at small effect sites, *θ*_*L*_ = 4*NLµ* is the total population scaled mutation rate of the trait.

We can then solve for the value of *δ*(*a*_*L*_) via a line search. First, we plug *T*^∗^ into the right hand side of Equation (D.4) and set it equal to the right hand side of Equation (D.8), yielding

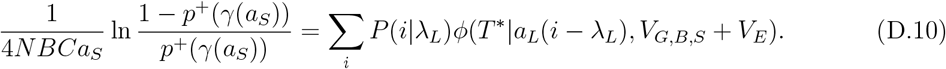

Then, for a proposed value of *δ*(*a*_*L*_), we compute *λ*_*L*_ using Equation (D.1), plug it into the RHS of Equation (D.10), and numerically solve for the value of *T*^∗^ that satisfies Equation (D.10). We can then conduct a line search for the value of *δ*(*a*_*L*_) that satisfies Equation (D.2), given the value of *T*^∗^ obtained from solving Equation (D.10).

To constrain the liability-scaled heritability to a specific value, we add one additional step. Given the proposed value of *δ*(*a*_*L*_), we compute the large effect contribution to the genetic variance as

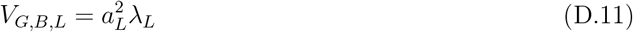

and then compute the environmental variance as

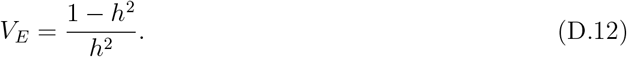

Note that the inclusion of the *B* value in Equations (D.8) and (D.9) to account for the impact of BGS on weakly selected sites represents the only substantial difference between the algorithm presented in Berg et al. (2025) and the one presented here. The solution in the case of no BGS is therefore obtained simply by setting *B* = 1. To compare cases with and without background selection, we first solve the above system of equations for a fixed value of *h*^2^ with *B* = 1, to obtain the solution in the absence of background selection. We then solve in the presence of background selection, holding the environmental variance (as opposed to the heritability), constant at the value we obtained in the no BGS case.

## E. An Incidental Finding That the Bulmer Effect Increases the Genetic Variance in the Long Term

In our multi-locus stabilizing selection simulations, when we used the same recombination rate between causal loci as used in the simulation of the liability threshold model (5 × 10^−6^), the genetic diversity reduction of causal sites under background selection initially did not match our expectations based on our theory and our single-locus simulations (Figure S11). We hypothesized that this might be related to the Bulmer effect (Bulmer 1971; Bulmer 1976), i.e. the accumulation of negative linkage disequilibrium between causal sites due to selection. More precisely, the genetic variance can be composed into two components:

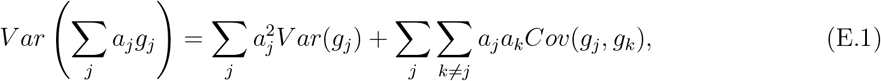

where the first term is the additive genic variance and the second term captures the contribution of linkage disequilibrium among causal sites. Under neutrality, the expectation of the LD term is equal to zero. However, because stabilizing selection acts most strongly against individuals carrying combinations of alleles that result in extreme phenotypes, it causes negative covariance among alleles of like sign, leading to a negative expectation for the contribution of the LD term. Thus, while stabilizing selection is acting, it leads to a reduction in the total genetic variance due to this negative LD. The reduction in genetic variance due to this negative LD is what is known as the “Bulmer effect”. Because recombination will quickly eliminate this negative LD if stabilizing selection ceases, the Bulmer effect is generally understood as a “short term” phenomenon.

For any given focal allele, the negative LD induced by the Bulmer effect leads to an attenuation of the allele’s marginal correlation with the phenotype, relative to its causal effect (Bulmer 1971; Veller and Coop 2024). In the modern statistical genetics literature, this effect is sometimes known as “linkage masking” (Brown et al. 2016). Because the expected change in the frequency of an allele from a given generation to the next depends on its marginal correlation with fitness in that generation, rather than its causal effect on fitness, the attenuation of the marginal phenotypic correlation in turn leads to an attenuation of the selection coefficient, and this in turn leads to a reduction in the magnitude of the expected change in frequency due to selection. More precisely, whereas in the absence of any Bulmer effect, the expected change in the minor allele frequency is given by:

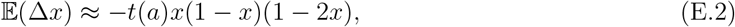

with 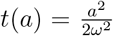, in its presence we expect that 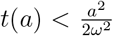, where the magnitude of the reduction in *t*(*a*) depends on the amount of negative LD. This slowdown in the pace of frequency change is studied in greater depth by Negm and Veller (2024).

This effect has an intriguing implication. That is, while the negative LD induced by stabilizing selection causes a short-term reduction in the genetic variance relative to the genic variance, over long timescales, we would expect it to increase in the genic variance, due to the weaker selection on individual variants. Whether the net effect is to increase or decrease the genetic variance then depends on which effect is larger.

We hypothesized that this phenomenon may be responsible for the difference that we initially observed between our predictions and our simulation results. To investigate this hypothesis, we performed simulations under the stabilizing selection model without background selection and varied the rate of recombination between the neighboring causal sites (Figure S11). When the recombination rate is low, we find that the genetic variance is reduced relative to the genic variance, but is nevertheless increased relative to what we observed with higher recombination rates (which match the theoretical predictions based on unlinked theory; Simons et al. (2018)). Thus, our simulation results suggest that over long time scales, the net impact of the Bulmer effect is to increase the genetic variance, as the increase in the genic variance caused by the slowdown in the pace of allele frequency change is larger than the reduction in variance due to the negative LD. Moreover, when we increased the recombination rate in our study of background selection to a level at which there was no longer any Bulmer effect or inflation of the genic variance, we found that our theoretical prediction for the impact of background selection then matched our theoretical predictions (Figure 4A), indicating that the Bulmer effect was indeed the culprit.

Our primary focus was on understanding the impact of background selection, so we did not study this effect any further. However, it is worth noting that, in addition to Negm and Veller (2024), at least two prior publications have also studied this effect. Specifically, Lande (1975) found that in his Gaussian allelic model, the decrease in genetic variance due to the Bulmer effect is exactly offset by the increase in genic variance, leading to no net effect. In contrast, Turelli and Barton (1990), employing their “rare alleles” approximation, do predict an increase in the genetic variance when recombination rates are low, but their predictions do not match our simulation results. A more complete analysis of this phenomenon would be valuable.

## F. Variation in a and B in the Threshold Model

In this section, we provide more explicit mathematical support for the arguments in the main text for the cases with variation in local *B* value. Here, we start from a slightly more general position than in the main text, by assuming that the effects and the local *N*_*e*_ reductions have some joint distribution *g*(*a, B*). Then analogous to equation (18) in the main text, equilibrium is established by evolving the value of *f* (*T*) that solves

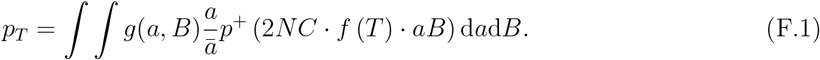

### Effectively neutral regime

When 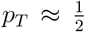, the fixation asymmetry is approximately linear, 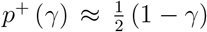. Inserting this approximation and evaluating the resulting integral, we have

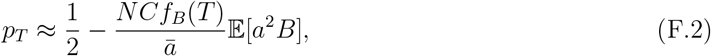

which we can solve for

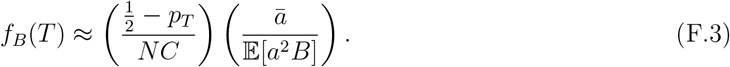

In the absence of background selection, *B* = 1, so

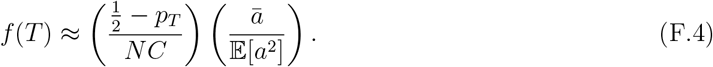

The global compensation factor is

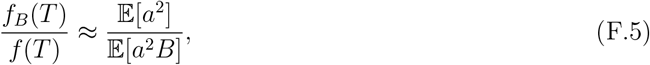

and is thus an average of *B* that is weighted by the squared effect sizes, reflecting the fact that variance contributions scale with *a*^2^ in the neutral regime. If *a* and *B* are independent, then

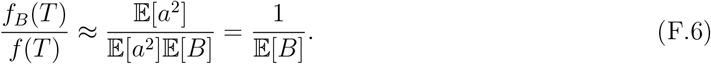

### Strong selection limit

In the strong selection limit, the behavior depends on whether the coefficient of variation of the effect distribution is large or small. Let us first consider the small coefficient of variation case.

### *g*_*a*_(*a*) has a small coefficient of variation

In this case, the distribution of scaled selection coefficients *γ* is centered over a relatively narrow range. Most sites will have similar scaled selection coefficients, and the bulk of their distribution moves from the weak into the strong selection regime as *p*_*T*_→ 0. In this regime, *p*^+^ (*γ*) = (1 + *e*^2*γ*^)^−1^ ≈ *e*^−2*γ*^, so equation (F.1) becomes

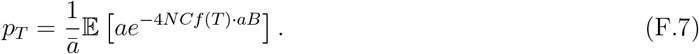

The exponential decay dominates over the linear factor of *a*, and so the primary contribution to the expectation is from sites with the minimum value of *aB*, i.e. the sites with the smallest effect sizes experiencing the strongest background selection. This implies that

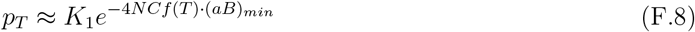

where *K*_1_ is a constant, and (*aB*)_*min*_ is the minimum of the product of *aB* across sites. Solving for *f* (*T*), we have

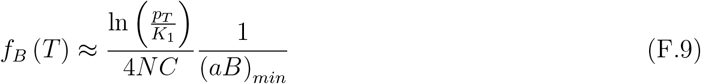

and

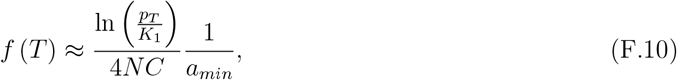

so that the global compensation factor is

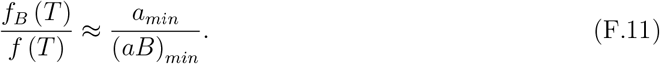

If *a* and *B* are independent, then (*aB*)_*min*_ = *a*_*min*_*B*_*min*_, and so

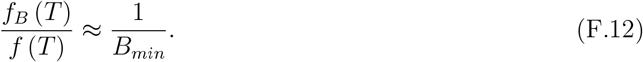

### *g*_*a*_(*a*) has a large CV

Alternatively, if there is high variance in the effect distribution, then the scaled selection coefficients of sites will not all cluster together in the same selection regime, but rather will be spread across all of the selection regimes, with a large subset of small effect sites belonging to the effectively neutral regime, even as *p*_*T*_ → 0. As a result, we must consider contributions from the whole distribution, rather than just a narrow slice of it. Here, it is not clear *a priori* how to obtain any general results for an arbitrary joint distribution, *g*(*a, B*), so we will assume they are independent (i.e., *g*(*a, B*) = *g*_*a*_(*a*)*g*_*B*_(*B*)). As a tractable choice for a distribution which can model the high variance scenario of interest, suppose that the effects follow a gamma distribution with shape and scale parameters *k* and *ω*, i.e. 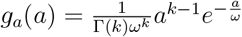 . The coefficient of variation of the gamma distribution is equal to *k*^− 1*/*2^, so we will be interested in the *k* → 0 limit, though we proceed generally for now, and consider this limit below.

Although the entire distribution contributes to the integral, the exponential suppression of the transition into the strongly selected regime still dominates, so we can still make the same *p*^+^ (*γ*) ≈ *e*^−2*γ*^ approximation that we made in the low CV case. Equation (F.1) becomes

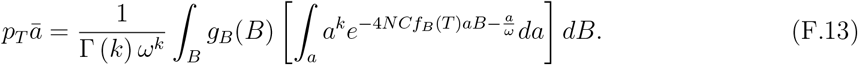

Because we are interested in the strong selection limit (i.e. *p*_*T*_ → 0), we assume that the threshold density *f*_*B*_ (*T*) is large enough that 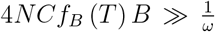 . In physical terms, this inequality implies that the fixation asymmetry *p*^+^ (*γ*) decays with increasing effect size *a* much more rapidly than the mutational input *g*_*a*_(*a*) does. That is, selection effectively prevents the fixation of deleterious alleles with moderate effect sizes well before such alleles become rare in the mutational distribution. Consequently, the integral is dominated by the interaction between selection and the power-law portion of the effect size distribution, allowing us to ignore the exponential suppression of the tail of the gamma distribution. Mathematically, this allows us to approximate the exponential function in equation (F.13): 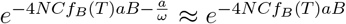, so that we can write

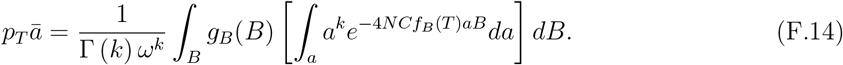

We can then use *u* substitution (set *u* = 4*NCf*_*B*_ (*T*) *aB* so that 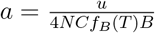 and 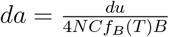) to rewrite the inner integral as

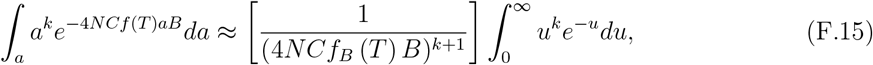

where the integral in *u* is the gamma function, i.e. 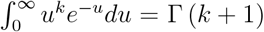. We can therefore write equation (F.14) as

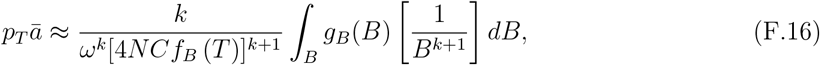

which we can solve to find

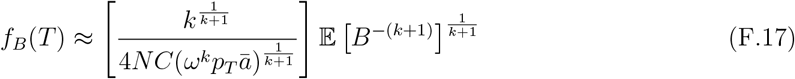

implying that

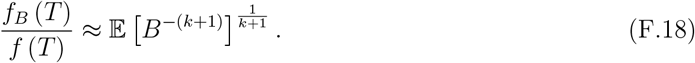

Now, let us consider the large coefficient of variation limit. As *k* → 0, the gamma distribution follows a power law that decays at rate *a*^−1^ until cutoff by the exponential suppress term 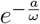 . In this case, the effect size distribution is very broad, and ^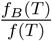^ is equal to the expectation of *B*^−1^:

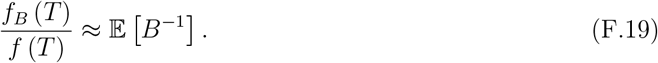

Alternatively, as *k* → ∞, the variance of the effect size distribution shrinks, and ^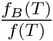^ converges toward the minimum,

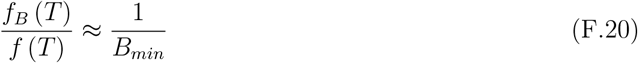

consistent with our previous results.

Finally, we must address the validity of the continuous approximation in the high-CV limit. In our continuous mathematical model, the Gamma distribution provides an effectively infinite number of sites with vanishingly small effect sizes. Consequently, the compensation process never terminates because there are always sites small enough to remain in the effectively neutral regime. In a physical genome with a finite number of sites *L*, there is necessarily a single site with the minimum effect size, *a*_*min*_. Technically, the compensation process described above would terminate once *f* (*T*) becomes large enough that even this smallest effect is pushed into the strong selection regime. However, this termination occurs only at the limit where selection is so strong that carrying a single allele at any site with any effect size is sufficient to cross the threshold (i.e., *s* ≈ *C*). Thus, the limit where this global compensation ultimately stops acting is precisely the limit where the epistasis becomes unimportant, and the model collapses onto one where all loci impact fitness independently in a Mendelian fashion. Thus, by invoking a model with threshold epistasis and wide variation in effect sizes in the context of a finite genome, we are effectively invoking the *p*_*T*_ → 0 limit, but assuming that *p*_*T*_ nonetheless remains large enough that *a*_*min*_ ≪ *p*_*T*_ . As shown in Figure 3B, explicit numerical solutions of equation (F.1) exhibit convergence to the *f*_*B*_ (*T*)*/f* (*T*) ≈ 𝔼 [*B*^−1^] limit at relatively large values of *p*_*T*_ once the CV becomes moderately large, so this assumption is entirely appropriate.

### Impact on the genetic variance

Having determined the global compensation factor, we can now derive the impact of background selection on the genetic variance contributed by specific sites. Let 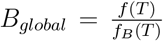 be the effective genome-wide reduction in the threshold density (note that this is the inverse of the compensation ratio derived above, e.g., 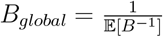 in the high-CV gamma limit).

Consider a specific site *i* with effect size *a* and a local effective population size reduction *B*_*local*_. In the absence of background selection, the site has a scaled selection coefficient *γ* = 2*Naf* (*T*)*C*. In the presence of background selection, the unscaled selection coefficient increases due to the global shift in threshold density:

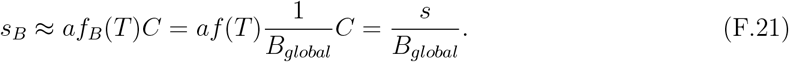

However, the local effective population size decreases: *N* → *NB*_*local*_. The new scaled selection coefficient is therefore:

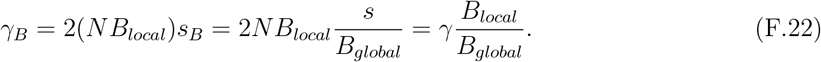

This ratio ^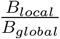 *f*^determines whether the site experiences a net increase or decrease in the efficiency of selection relative to drift.

The genetic variance contributed by this site is proportional to the mutation rate and the function *b*(*γ*)*/γ*, where *b*(*γ*) = tanh(*γ*). Accounting for the reduction in the effective mutation rate (*θ* → *θB*_*local*_), the variance in the presence of background selection is:

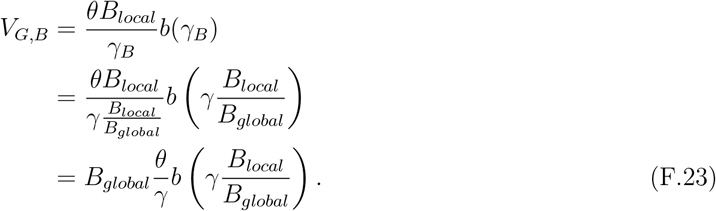

Comparing this to the original variance 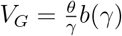, we obtain the reduction factor:

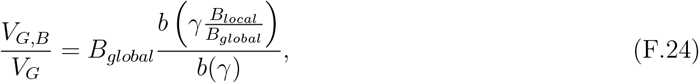

i.e. equation (22) in the main text.

## Supplementary

**Figure S1:**
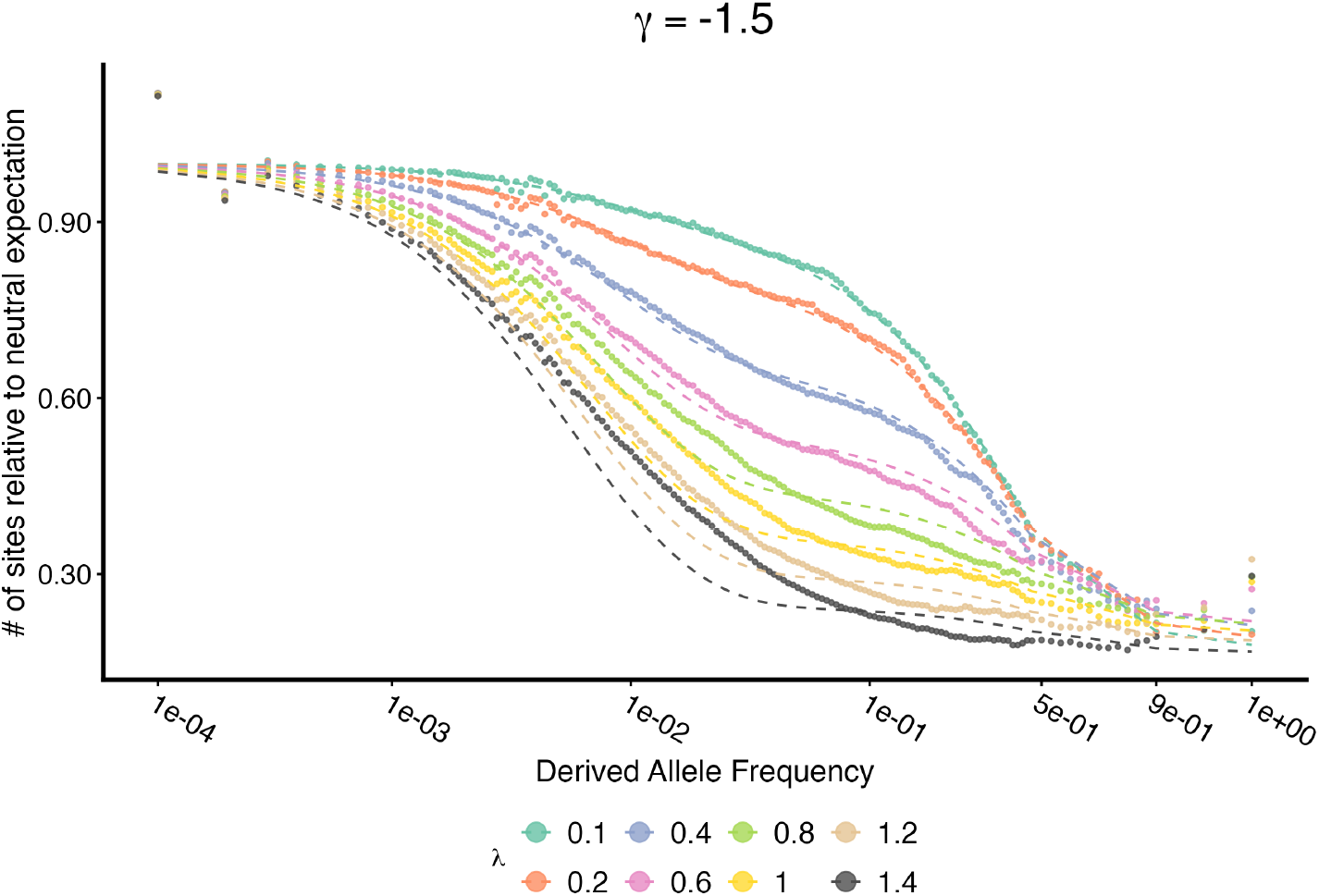
Robustness of the frequency-dependent BGS approximation across varying mutation intensities (*λ*). The site frequency spectrum (SFS) of a focal site with *γ* = −1.5, scaled relative to the standard neutral expectation, is shown for simulations with varying values of the background mutation flux *λ* = 2*ν/r*. Points represent simulation results, while solid lines show the analytical prediction using the *B*(*q*) approximation derived in the weak mutation limit (Equation (A.27)). Although the analytical model is formally derived for the limit where mutation is weak relative to recombination (*λ* ≪ 1), it accurately captures the distortion of the SFS for *λ <* 1*/*2. Departures from the prediction for *λ >* 1*/*2 indicate the onset of the “fitness drag” regime driven by the accumulation of new mutations, which we describe in Supplementary Text A.

**Figure S2:**
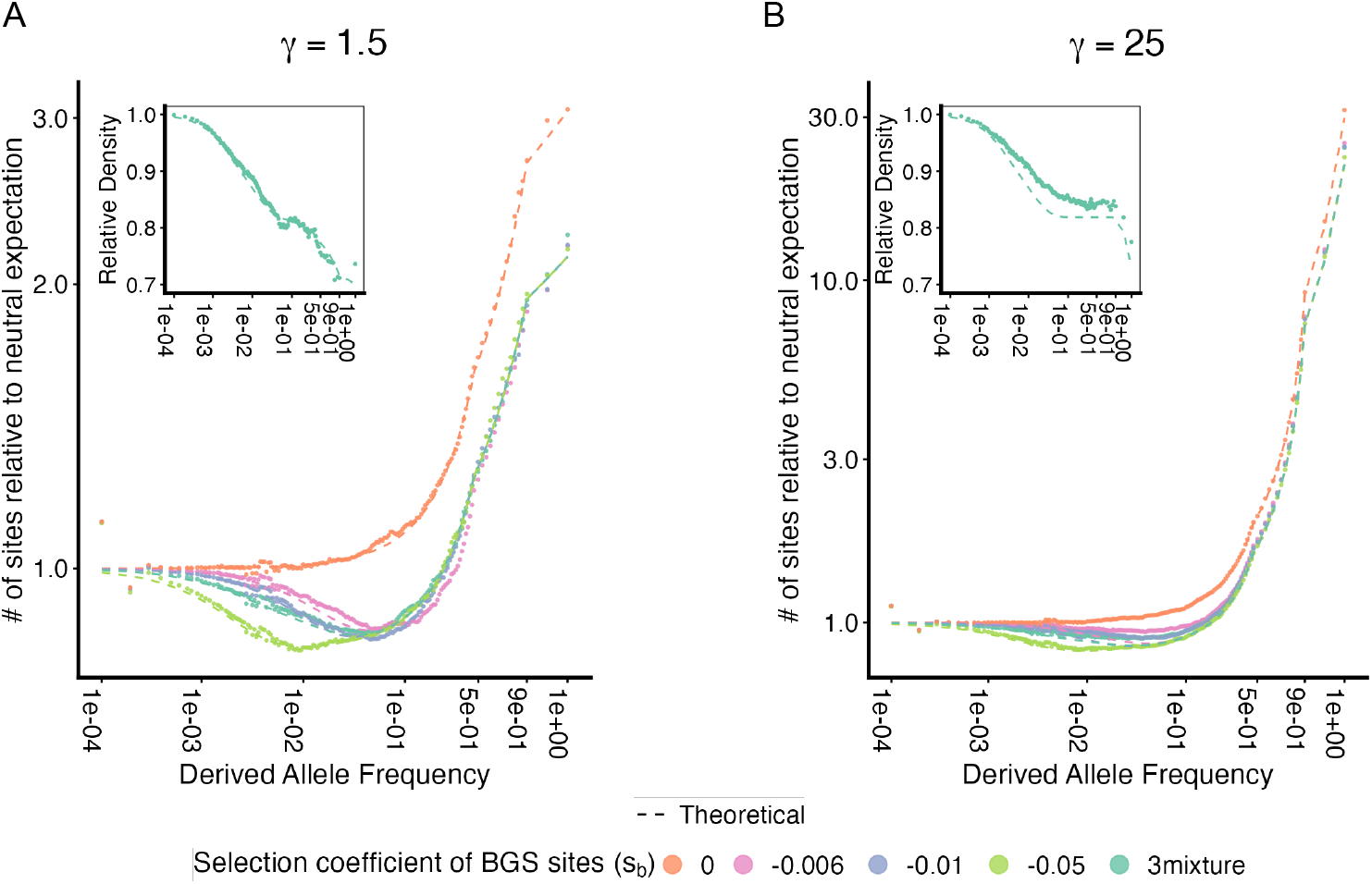
The site frequency spectrum (SFS) for positively selected focal alleles, scaled to the neutral allele expectation, is shown for a BGS intensity of *B ≈* 0.82. Points represent simulation results, while dashed lines indicate theoretical predictions based on the frequency-dependent *B*(*q*) approximation (Equation A.32). We show results for three different selection coefficients of background selection sites (*s*_*b*_ = −0.006, −0.01, −0.05), as well as an equal mixture of all three. The insets show the relative density of alleles across frequency bins, comparing scenarios with BGS to those without. Focal selected alleles are under (A) weak positive selection (*γ* = 1.5) and (B) intermediate positive selection (*γ* = 25). When *γ* = 25, simulations with weaker BGS (|*s*_*b*_| < 0.05) exhibit a deviate from the theoretical line. This occurs because when the strength of direct selection on the focal allele is similar to the strength of background selection ( |*γ*| *≈* |2*Ns*_*b*_|), the two processes act on similar timescales. This transits into the regime where the focal allele is dominated by its own fitness effect before the background is purged (Equation A.35).

**Figure S3:**
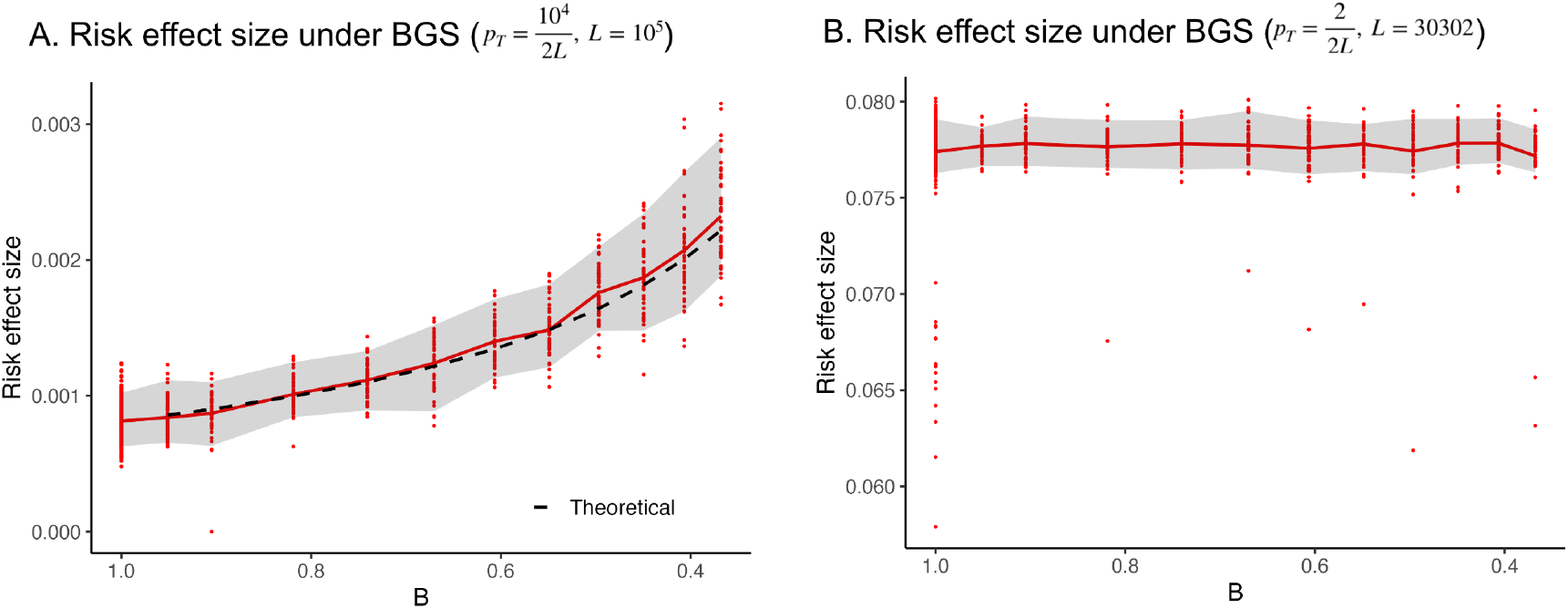
The impact of BGS on risk effect size in the single-effect liability threshold model. (A) When 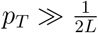, the risk effect size increases with increasing strength of BGS. The black dashed line shows the theoretical prediction of *f* (*T*) → *f* (*T*)*/B*. (B) When 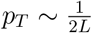, the risk effect size does not change under BGS.

**Figure S4:**
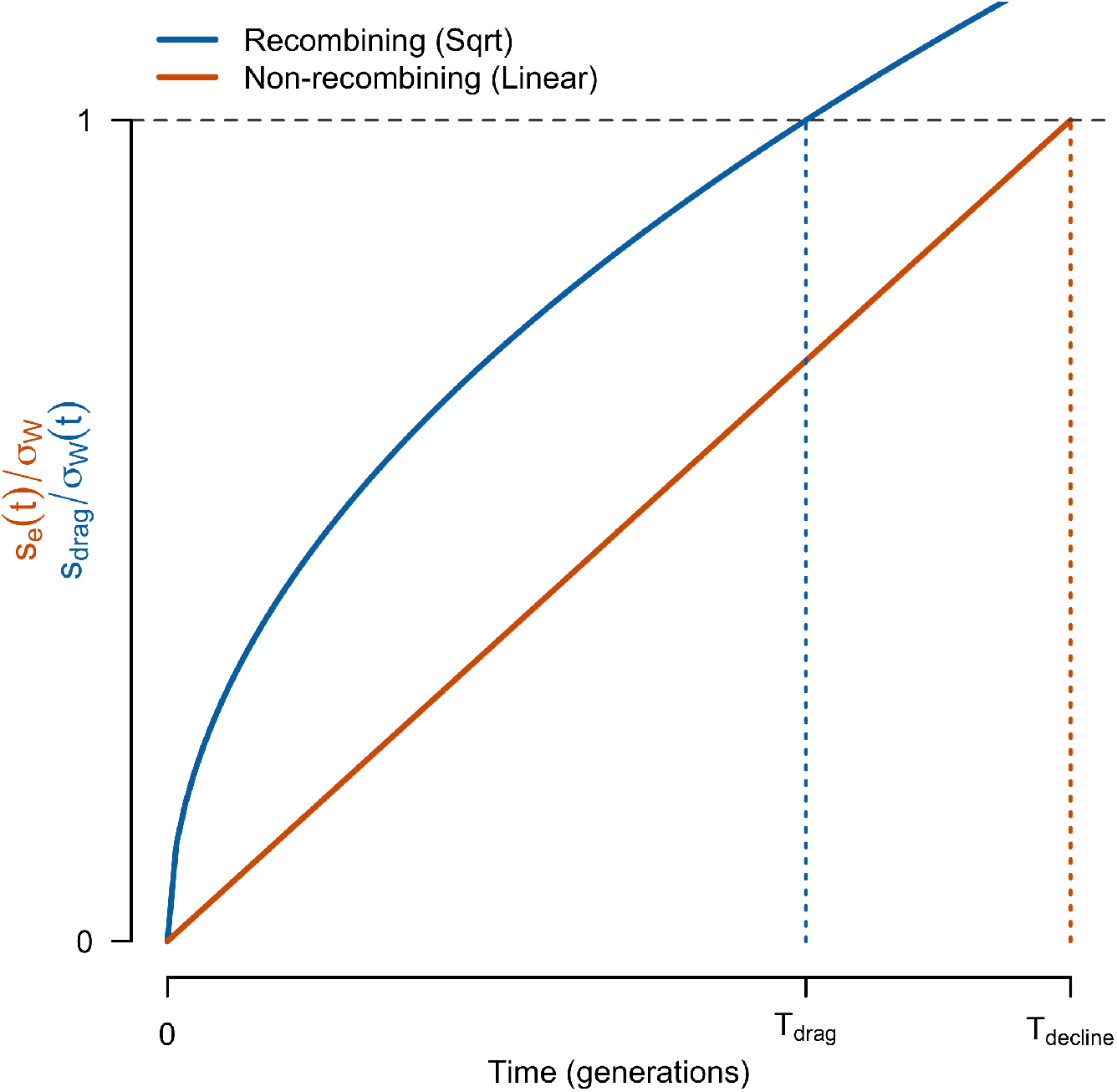
Dynamics of the onset of effective selection in recombining vs. non-recombining models. This figure compares the time evolution of the ratio of the effective selection coefficient (*s*_*eff*_) to the relevant standard deviation of fitness (*σ*_*W*_) for a newly arisen allele. In the non-recombining model (orange), the fitness deficit accumulates linearly (*s*_*eff*_ (*t*) ∝ *t*) against a constant background variation, leading to a linear increase in the ratio. The threshold for effective selection (*T*_*decline*_) is reached when this ratio equals 1. In the recombining model (blue), the fitness deficit is constant (*s*_*drag*_ = *λs*_*b*_), but the relevant background variation shrinks as the associated haplotype block shortens (*σ*_*W*_ (*t*) ∝ *t*^−1*/*2^), causing the ratio to grow with 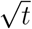. This dynamic leads to a more rapid onset of effective selection, both because *T*_*drag*_ *< T*_*decline*_ and because the functional form of the ratio results in faster growth of the relative fitness deficit in early generations. Together, these factors imply that the frequency spectrum is distorted at significantly lower frequencies in recombining genomes. For the example plotted here, *λ* = 2.

**Figure S5:**
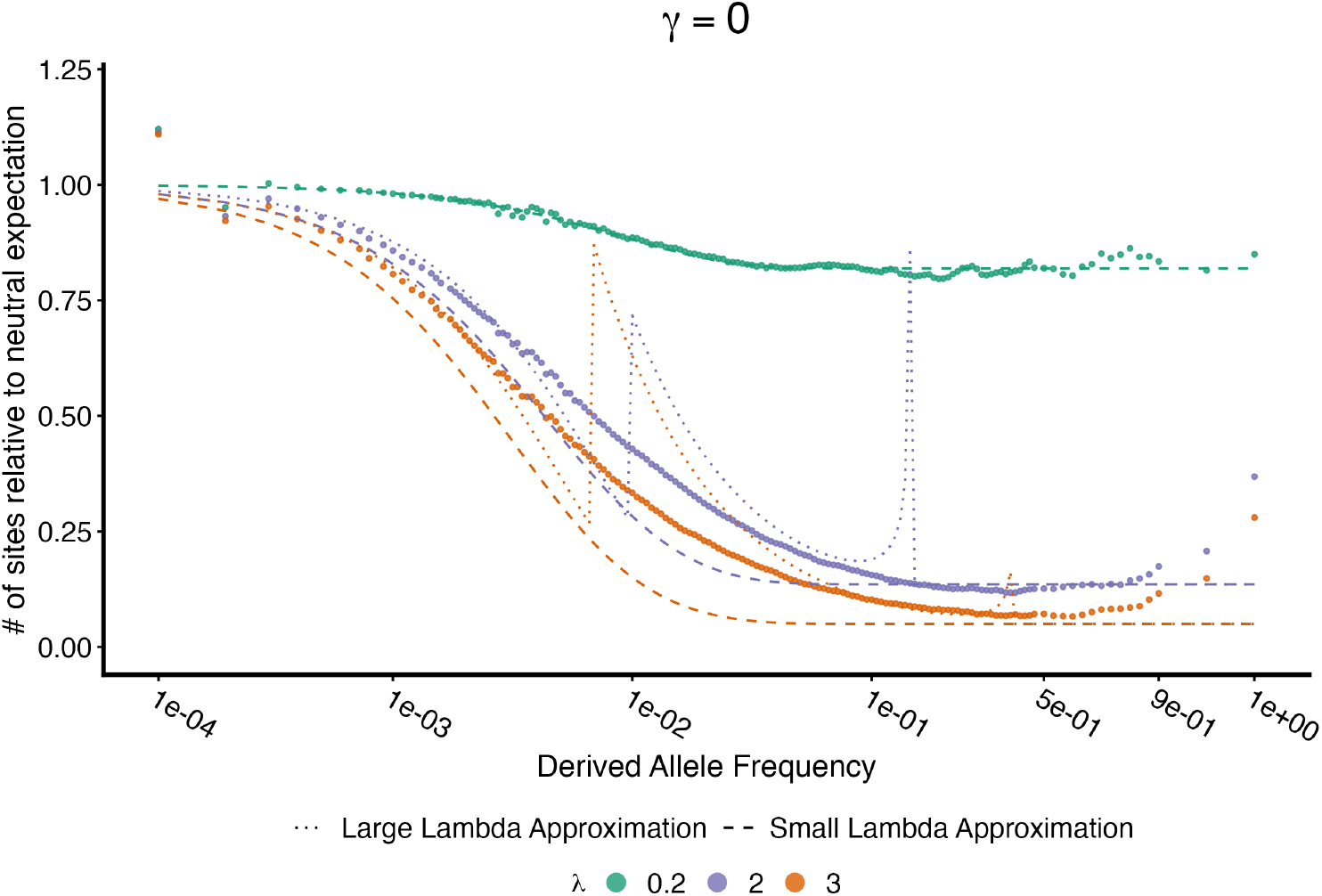
The site frequency spectrum (SFS) of a neutral focal site (*γ* = 0) is shown relative to the standard neutral expectation for three different background selection intensities. Points represent simulation results for various levels of background mutation flux (*λ* = 2*ν/r*), ranging from weak (*λ* = 0.2) to strong (*λ* = 2, 3). For low mutation flux (*λ* = 0.2), the distortion of the SFS is well-captured by the weak mutation/strong recombination (small lambda) approximation with Equation A.24 (dashed lines), which accounts for selection acting primarily against deleterious alleles already present on the initial background. As the mutation flux increases (*λ >* 1*/*2), “fitness drag” regime kicks in and the dynamics are instead better described by the strong mutation/weak recombination (large lambda) approximation with (Equation A.47)(dotted lines), which accounts for the continuous accumulation of fitness drag on the shortening associated haplotype block. Notably, the spikes in the large lambda approximation represent logarithmic divergences near the transitions between frequency regimes in Equation A.47, where Cvijović et al. (2018) applied a smoothing technique to resolve these boundaries.

**Figure S6:**
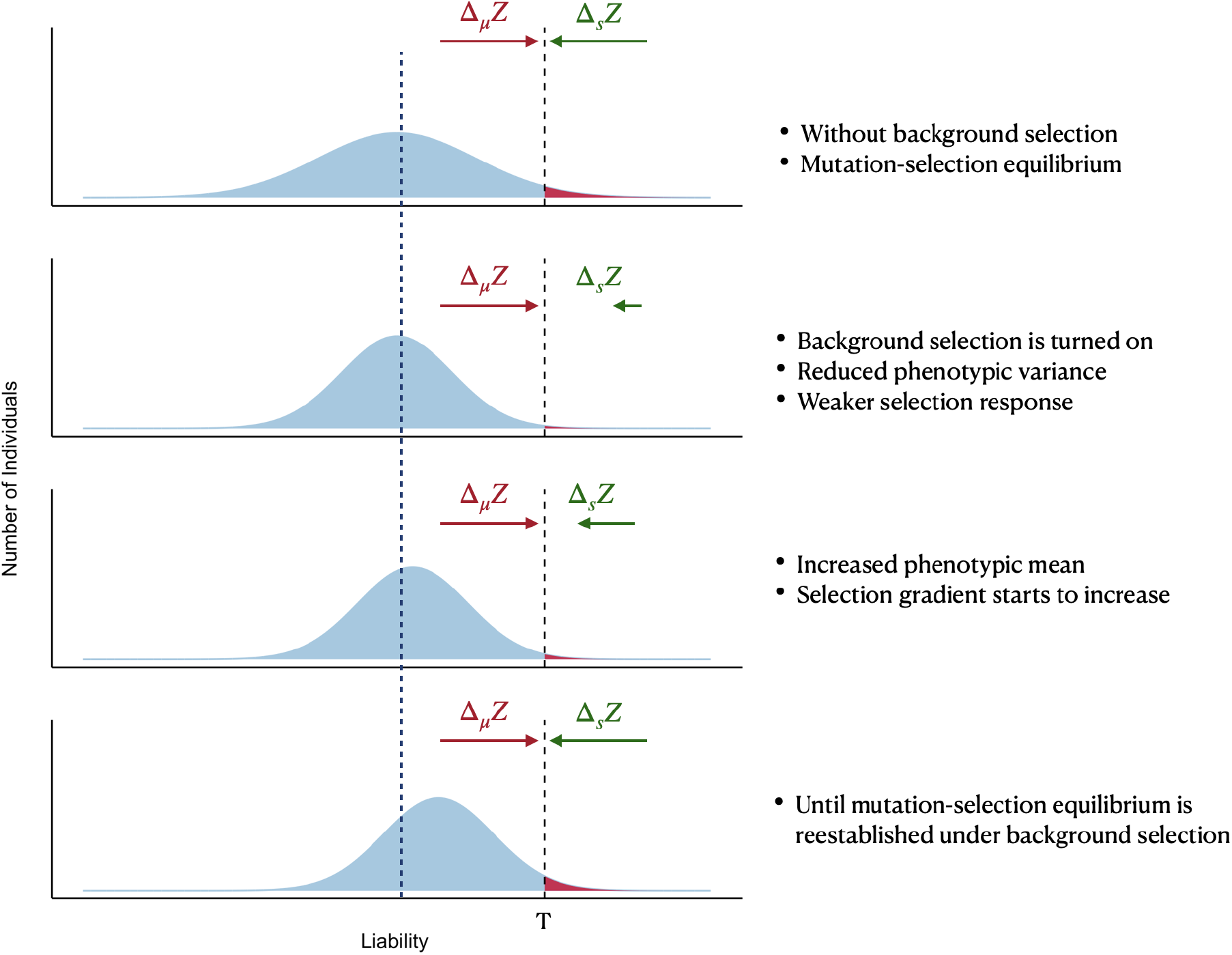
A cartoon illustration that explains how background selection impacts disease prevalence. This illustration imagines a dynamic process where the population is first at equilibrium in the absence of background selection, before it is “turned on”, and the population evolves to the new equilibrium. In reality, top and bottom panels should be viewed as counterfactual comparison in a model without vs with background selection.

**Figure S7:**
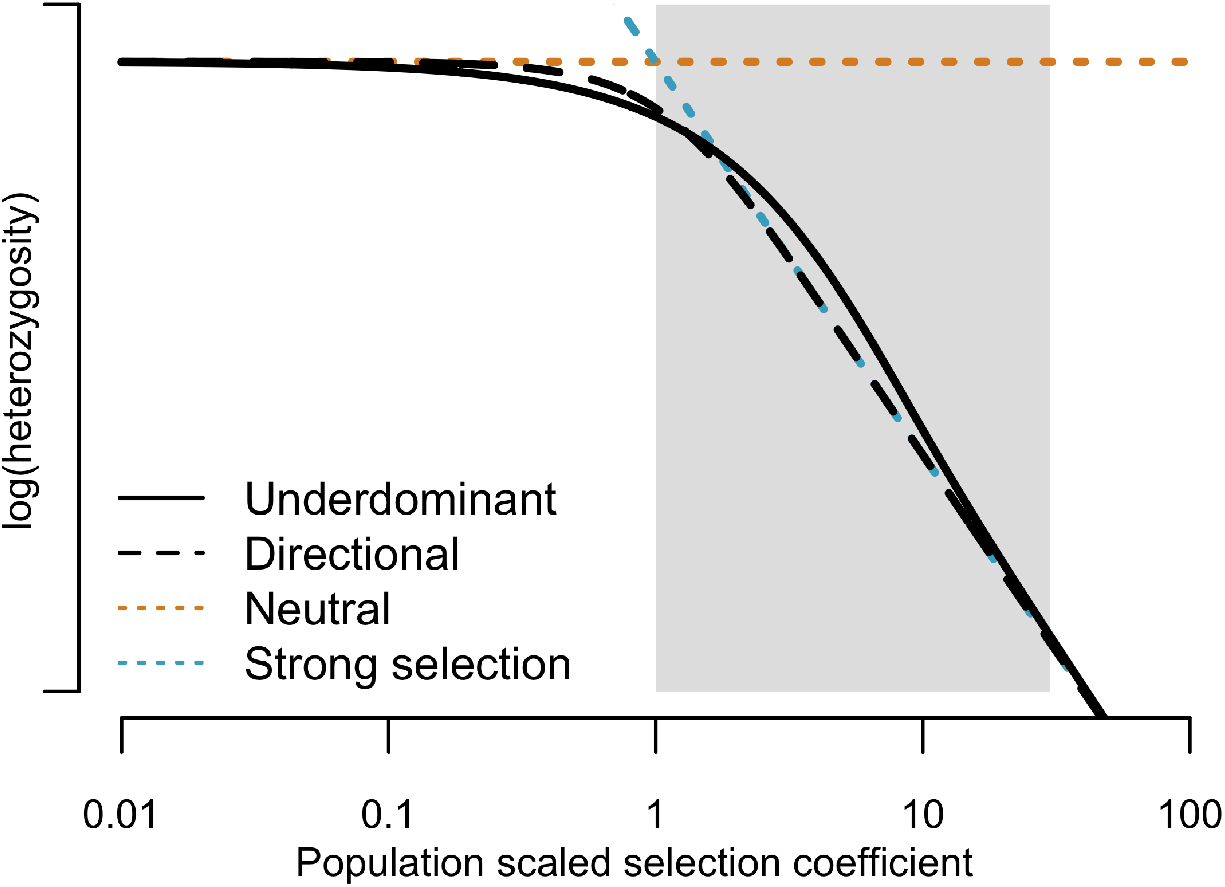
A comparison of expected heterozygosity for alleles with directional selection (Equation (6), dashed black line) vs. under dominant selection (Equation (23), solid black line) at different scaled selection coefficients. The expected neutral heterozygosity is *θ*, and the expected heterozygosity for strong selection is ^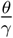^ .

**Figure S8:**
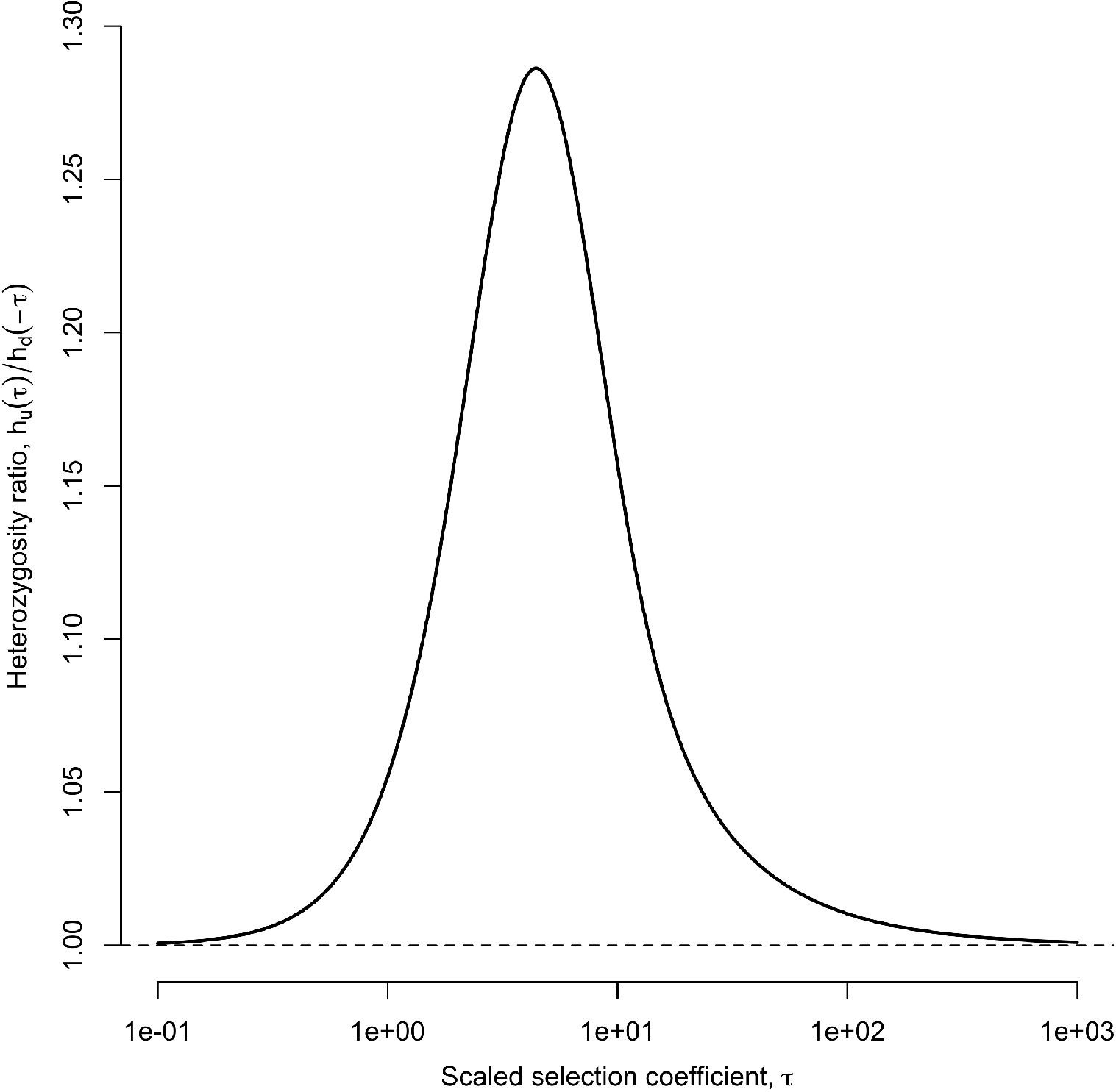
The ratio 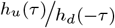, measuring the increase in heterozygosity due to the underdominance-induced slow down in allele frequency change, as a function of *τ* .

**Figure S9:**
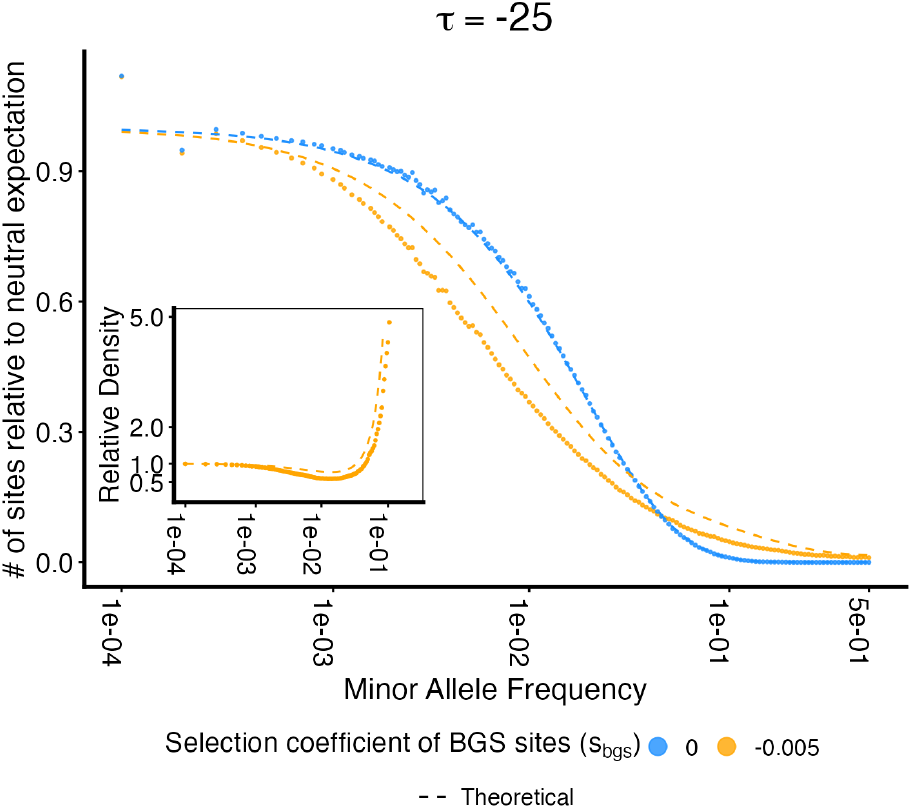
Site frequency spectrum (upper panel) and relative density (lower panel) of a selected allele with under-dominant selection 2*Nt* = −25, with a background selection intensity *B≈* 0.2.The theoretical results (dashed line) obtained with frequency dependent B approximation using equation A.26 breaks down in this large mutation/small recombination regime and does not predict the simulation well.

**Figure S10:**
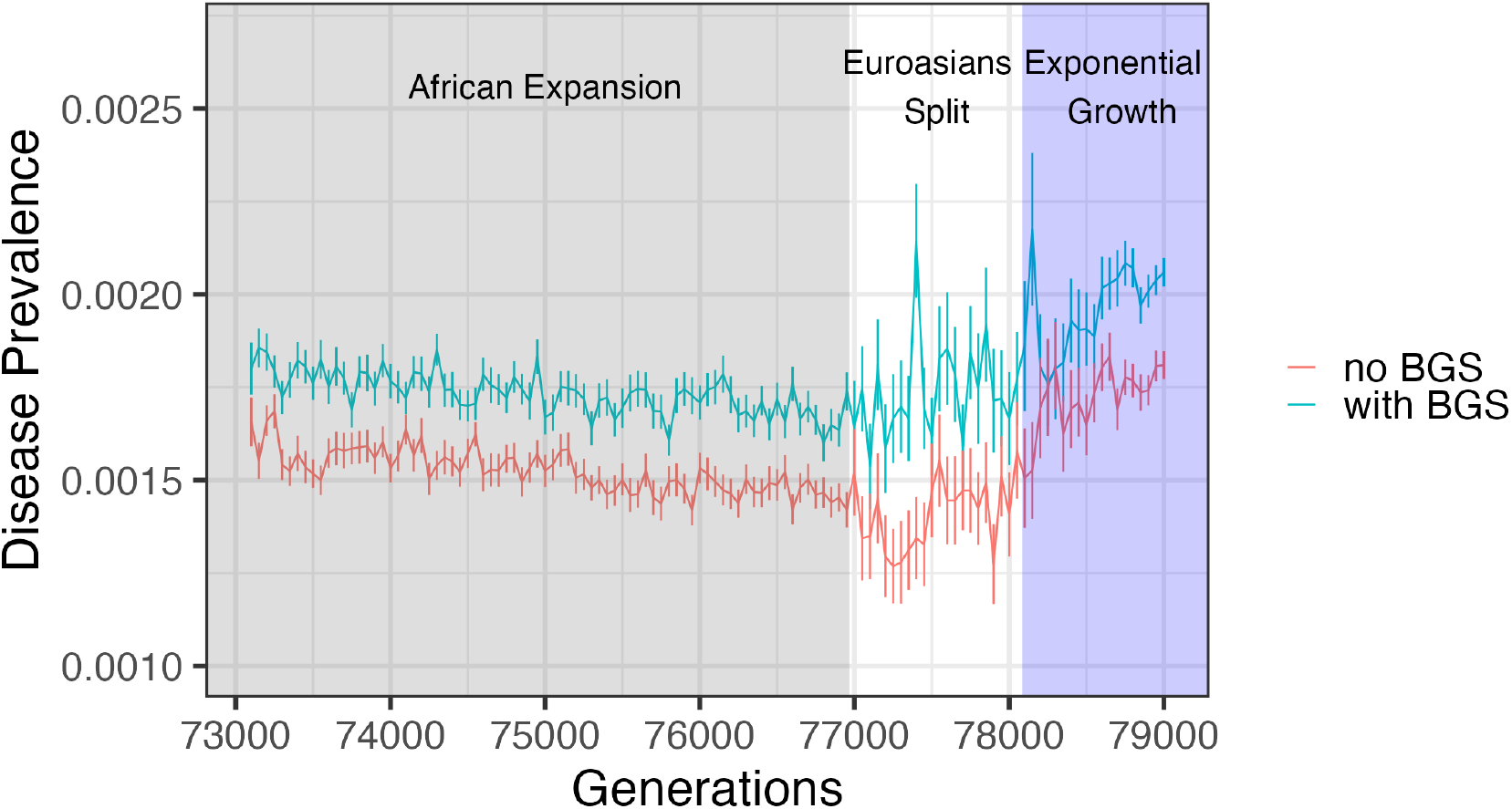
We simulated a “CEU-like” demographic history with the out-of-Africa demographic history (Gravel et al. 2011) with the strength of background selection set to reduce neutral variance by an average 17%, as estimated in humans (Murphy et al. 2022). We simulated a disease under the liability threshold model with heritability *h*^2^ = 0.5, and a mutational asymmetry *p*^+^(*a*) = 0.05. We reported the disease prevalence of the African population before the Euroasians split and of the European population after the split. We compared the disease prevalence with no background selection (red) and with background selection (green) intensity *B≈* 0.83 (average estimated in humans). The error bar represents the standard error of the mean among 100 replications. In these simulations, the prevalence of a polygenic disease with heritability 0.5 and weakly selected causal loci in the presence day (generation 79000) in the absence of background selection was 0.18% (95% CI: 0.176% - 0.191%), and its presence 0.21% (95% CI: 0.203% - 0.219%), an average increase of 14.9%.

**Figure S11:**
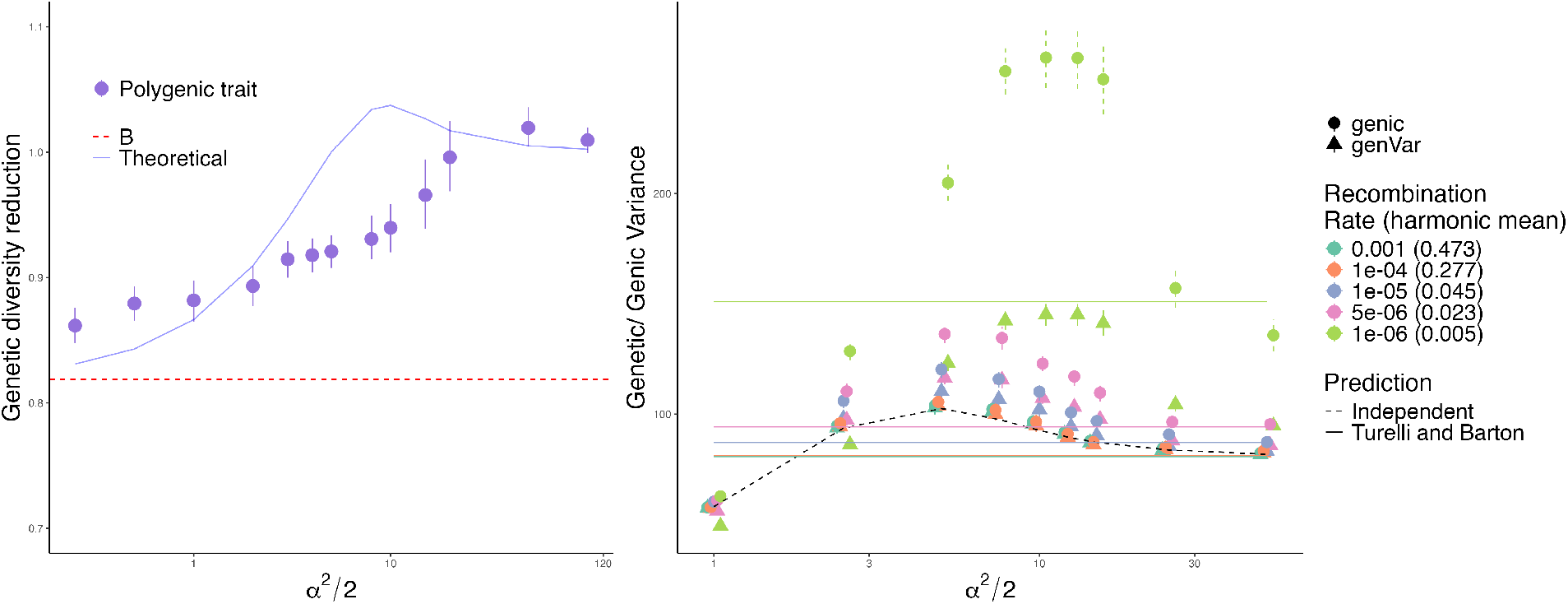
Left panel: we simulated polygenic traits evolving under stabilizing selection and set the recombination rate between causal loci at 5 × 10^−6^ and mutation rate at 2× 10^−8^. With these parameters choices, we found that the effect of BGS on the genetic variance of the trait was different from the single-site prediction. Right panel: we simulated traits under stabilizing selection without background selection and varied the recombination rates between causal loci. The mutation rate is fixed at 2 × 10^−8^. We plot the genic variance (circle) and genetic variance (triangle) and compare them to the predicted genetic variance that assumes independent causal loci (dashed line), and to the “rare alleles” approximation from Turelli and Barton 1990 (solid lines). While the LD component of genetic variance is negative, as expected, due to the slow down mechanism described by Negm and Veller (2024) the genic variance is increased by more than the negative LD subtracts, so that the net effect on the genetic variance is to increase it.

**Figure S12:**
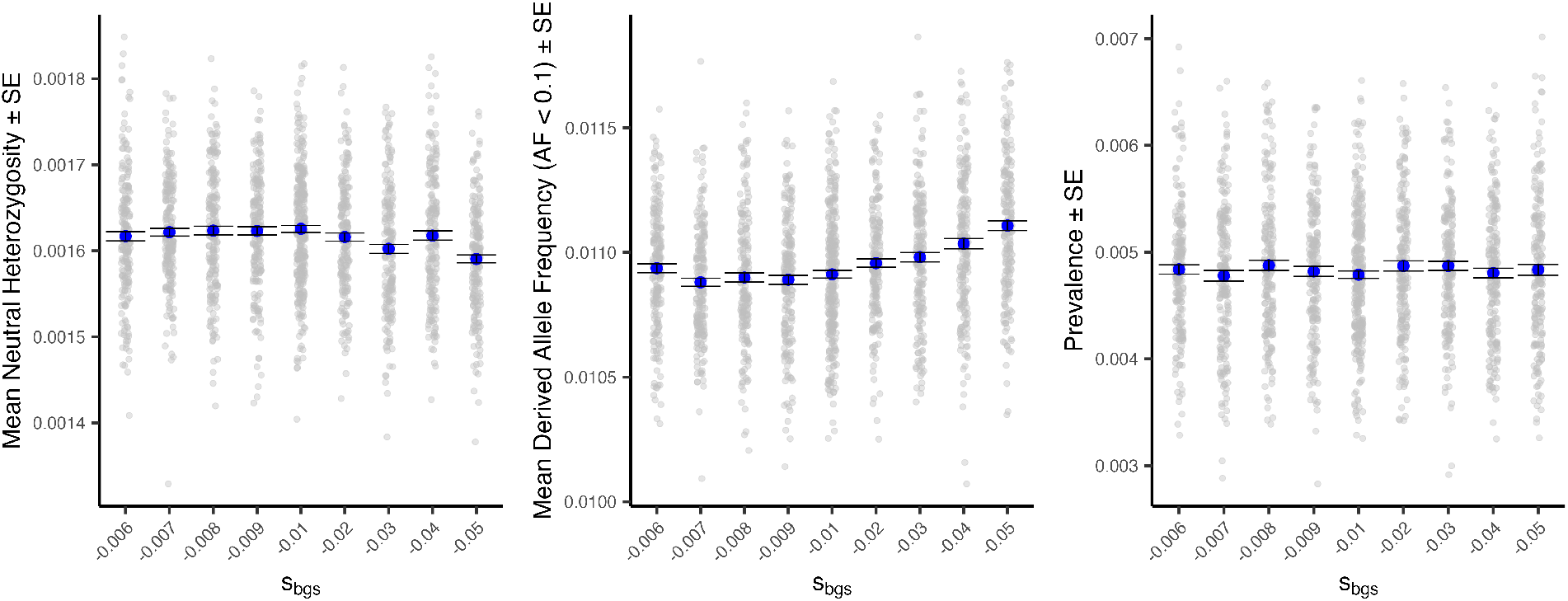
Simulations were designed to isolate the effect of background selection–induced distortion of the site-frequency spectrum (SFS), particularly the excess of rare variants, on disease prevalence under the liability-threshold model. Mutation rate (*ν* = 2 × 10^−8^) and recombination rate (*r* = 2 × 10^−7^) at BGS sites were held fixed, while only the selection coefficient (*s*_*b*_) was varied. The left panel shows that mean neutral heterozygosity is insensitive to *s*_*b*_. The center panel shows that weakening selection at BGS sites leads to a decrease in the mean derived-allele frequency among rare variants. Despite this shift in the SFS, the disease prevalence is not minimally affected, as shown in the right panel.

## Notes

### Competing Interest Statement

The authors have declared no competing interest.

### Summary of Updates

Several new results have been derived and the manuscript has been substantially rewritten.

